# The Prolyl-tRNA Synthetase Inhibitor Halofuginone Inhibits SARS-CoV-2 Infection

**DOI:** 10.1101/2021.03.22.436522

**Authors:** Daniel R. Sandoval, Thomas Mandel Clausen, Chelsea Nora, Adam P. Cribbs, Andrea Denardo, Alex E. Clark, Aaron F. Garretson, Joanna K.C. Coker, Anoop Narayanan, Sydney A. Majowicz, Martin Philpott, Catrine Johansson, James E. Dunford, Charlotte B. Spliid, Gregory J. Golden, N. Connor Payne, Mark A. Tye, Cameron J. Nowell, Eric R. Griffis, Ann Piermatteo, Kaare V. Grunddal, Thibault Alle, Jason A. Magida, Blake M. Hauser, Jared Feldman, Timothy M. Caradonna, Yuan Pu, Xin Yin, Rachael N. McVicar, Elizabeth M. Kwong, Ryan J. Weiss, Michael Downes, Sotirios Tsimikas, Aaron G. Smidt, Carlo Ballatore, Karsten Zengler, Ron M. Evans, Sumit K. Chanda, Ben A. Croker, Sandra L. Leibel, Joyce Jose, Ralph Mazitschek, Udo Oppermann, Jeffrey D. Esko, Aaron F. Carlin, Philip L.S.M. Gordts

## Abstract

**Summary Paragraph:** We identify the prolyl-tRNA synthetase (PRS) inhibitor halofuginone^1^, a compound in clinical trials for anti-fibrotic and anti-inflammatory applications^2^, as a potent inhibitor of SARS-CoV-2 infection and replication. The interaction of SARS-CoV-2 spike protein with cell surface heparan sulfate (HS) promotes viral entry^3^. We find that halofuginone reduces HS biosynthesis, thereby reducing spike protein binding, SARS-CoV-2 pseudotyped virus, and authentic SARS-CoV-2 infection. Halofuginone also potently suppresses SARS-CoV-2 replication post-entry and is 1,000-fold more potent than Remdesivir^4^. Inhibition of HS biosynthesis and SARS-CoV-2 infection depends on specific inhibition of PRS, possibly due to translational suppression of proline-rich proteins. We find that pp1a and pp1ab polyproteins of SARS-CoV-2, as well as several HS proteoglycans, are proline-rich, which may make them particularly vulnerable to halofuginone’s translational suppression. Halofuginone is orally bioavailable, has been evaluated in a phase I clinical trial in humans and distributes to SARS-CoV-2 target organs, including the lung, making it a near-term clinical trial candidate for the treatment of COVID-19.

## MAIN TEXT

Severe acute respiratory syndrome-related coronavirus 2 (SARS-CoV-2), the causative agent of coronavirus disease 2019 (COVID-19), has swept across the globe causing 32.1 million infections and 980,339 confirmed deaths as of September 24^th^ 2020^5^. In May 2020, the US Food and Drug Administration (FDA) granted an Emergency Use Authorization (EUA) for Remdesivir (RDV, GS-5734) for the treatment of hospitalized patients with severe COVID-19^6^. In the preliminary report of the ACTT-1 clinical trial subgroup analysis, Remdesivir significantly reduced time to recovery only in patients who were on low-flow oxygen at baseline^4^. This finding suggested that Remdesivir treatment in COVID-19 may provide greater benefit if started before the development of severe disease^7,8^. However, data from multiple trials suggests that Remdesivir provides modest clinical benefit compared with standard of care^4,9,10^. Furthermore, Remdesivir must be administered intravenously, which functionally prevents its use in pre-symptomatic or early symptomatic mild disease. Convalescent plasma from individuals who have recovered from COVID-19 has also been granted EUA for hospitalized patients with COVID-19 but efficacy data from randomized trials are needed. Hence, there is an urgent unmet clinical need for antiviral therapeutics that can reduce COVID-19 associated morbidity and mortality and that optimally could be administered early after symptom onset to prevent the development of severe respiratory disease^11,12^.

SARS-CoV-2 entry into lung upper airway epithelial cells depends on ACE2, TMPRSS2, and cell surface heparan sulfate (HS)^3,13,14^. Blocking the interaction of cell surface HS and spike protein attenuate SARS-CoV-2 binding and SARS-CoV-2 infection^3^. In this report we identified halofuginone, a synthetic analog of the natural product febrifugine that is derived from *Dichroa febrifuga* (one of the 50 fundamental herbs of traditional Chinese medicine), as a potent inhibitor of HS biosynthesis and SARS-CoV-2 infection^1,15^. We demonstrate that halofuginone reduces HS biosynthesis, thereby decreasing SARS-CoV-2 spike protein binding to cells, infection with SARS-CoV-2 spike protein pseudotyped VSV virus, and infection of authentic SARS-CoV-2. Additionally, halofuginone inhibited authentic SARS-CoV-2 replication post-entry. Inhibition of prolyl-tRNA synthetase (PRS) activity was responsible for both the HS-dependent and the HS-independent antiviral properties of halofuginone. These findings identify halofuginone as a translational regulator of cell surface HS biosynthesis and a potentially potent, orally bioavailable, therapeutic for the treatment and prevention of COVID-19.

## RESULTS

### Halofuginone Reduces SARS-CoV-2 Spike Protein Binding and Infection by SARS-CoV-2 Spike Protein Pseudotyped Virus

To identify potential agents that attenuate SARS-CoV-2 spike protein binding, we screened a library of small-molecule antagonists of host epigenetic regulators and protein translation elements of the prolyl-tRNA synthetase complex, along with their chemically matched inactive analogs^16–18^. The compound library has been successfully used to investigate targets mediating anti-inflammatory or anti-proliferative effects in a variety of biological contexts^16–18^ and importantly, contains several FDA-approved drugs, which may facilitate rapid deployment for treatment of COVID-19 patients. Hep3B human hepatoma cells were treated with the compound library at the indicated concentrations (Fig. 1a & **Extended data Fig. 1**) for 18 h, and binding of SARS-CoV-2 spike protein was assessed using recombinant receptor binding domain (RBD; from isolate Wuhan 20) in flow cytometry (Fig. 1a). As a positive control cells were treated with a mixture of heparin lyases, which digest HS chains^3^. Class 1 histone deacetylase inhibitors, such as Romidepsin (1 µM) or Belinostat (5 µM), decreased SARS-CoV-2 RBD binding by ∼50% compared to excipient control (DMSO; Fig 1a). Halofuginone (1 µM) reduced binding by 84%, a level of reduction similar to the impact of heparin lyase treatment (Fig 1a). Halofuginone targets the prolyl-tRNA synthetase (PRS) active site of the human glutamyl-prolyl tRNA synthetase (EPRS) and has been used in clinical and preclinical studies to treat fibrotic disease and to attenuate hyperinflammation^1,19^. We then treated Vero E6 African green monkey kidney epithelial cells, Caco-2 human epithelial colorectal adenocarcinoma cells, and Calu-3 human epithelial lung adenocarcinoma cells with halofuginone. Halofuginone reduced RBD protein binding to Hep3B, Calu-3 and Caco-2 but showed very modest effects in Vero E6 cells (Fig. 1b-e). The degree of inhibition was greatest in the Calu-3 line (5-fold reduction in RBD binding; Fig. 1e). To test the effect of halofuginone on the binding of spike protein in a more native presentation, we examined the ability of halofuginone to inhibit infection of SARS-CoV-2 spike protein pseudotyped VSV in Hep3B cells. Halofuginone inhibited infection by up to ∼30-fold in a dose-dependent manner (Fig. 1f). Consistent with the inability of halofuginone to reduce RBD binding to Vero E6 cells (Fig. 1c), no inhibition of infection was seen in these cells (Fig. 1g). We next tested if halofuginone inhibits the infection of authentic SARS-CoV-2 in Hep3B. Cells were treated with halofuginone at different doses for 24 h prior to and during infection (MOI of 0.1) (Fig. 1h)^20^. The culture medium was collected, and the presence of virus was measured by plaque assays in Vero E6 cells. No infectious SARS-CoV-2 could be detected in the supernatants of Hep3B cells treated with halofuginone at doses greater than 50 nM (Fig. 1h). To explore the effect of halofuginone on SARS-CoV-2 infection in a more clinically relevant model, we grew primary human bronchial epithelial cells grown at an air-liquid interface and infected them with authentic SARS-CoV-2 virus with and without halofuginone treatment. Halofuginone significantly reduced the number of SARS-CoV-2 infected cells at both 10 nM and 100 nM without affecting cell viability (Fig. 1i-j & **Extended data Fig. 2**).

**Figure 1.**
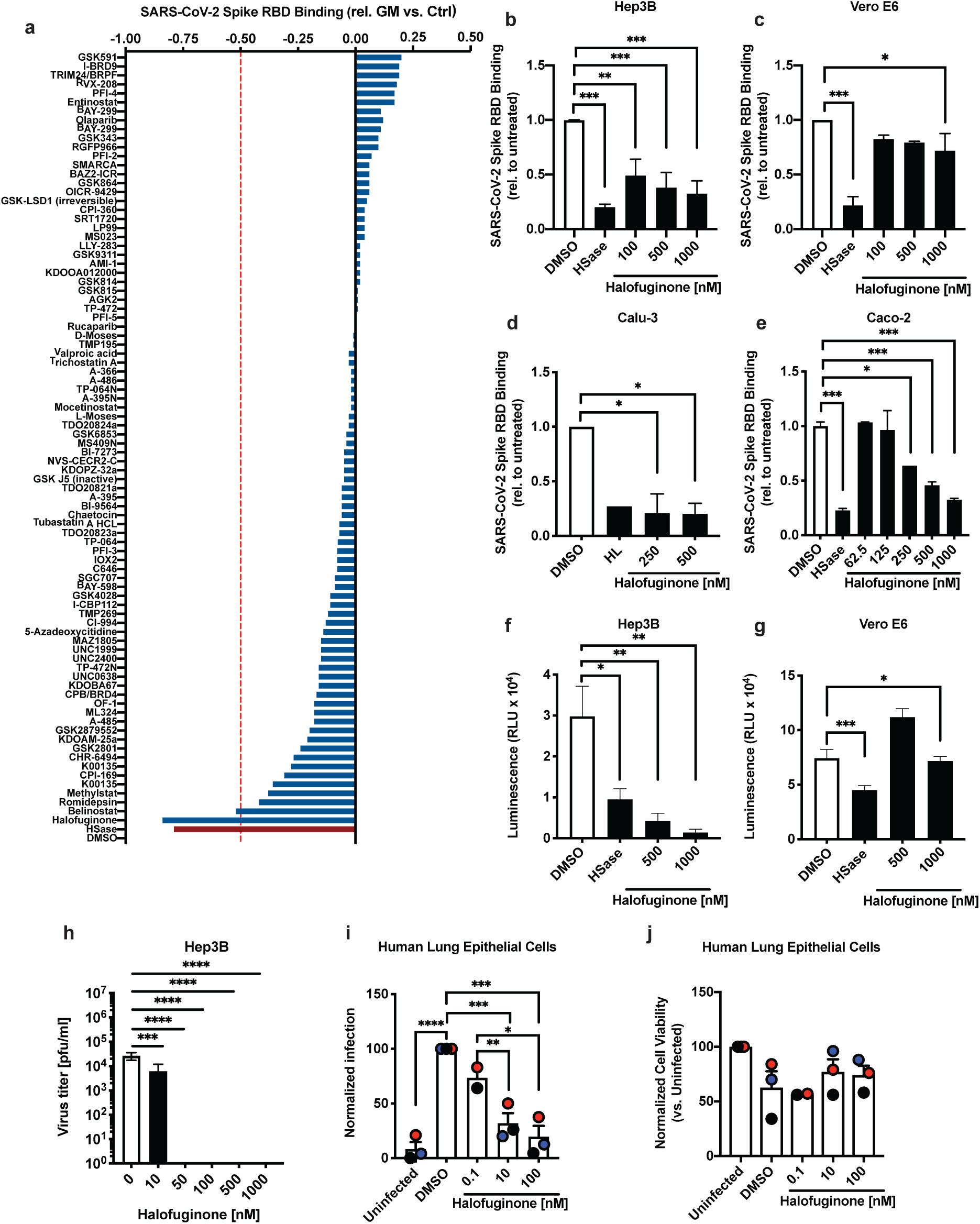
Screen of epigenetic and translational regulatory compounds identify Halofuginone as a potent inhibitor of SARS-CoV-2 Spike HS dependent cellular adhesion. **a,** Hep3B cells were treated with a library of epigenetic and translational regulatory compounds or heparin lyase (HSase) as a positive control and tested for their interaction with recombinant SARS-CoV-2 RBD protein. **b-e,** Titration of halofuginone on Hep3B cells (**b**), Vero E6 (**c**), Calu-3 (**d**) and Cacu-2 (**e**) cells (n= 2-4 replicates per condition), and its effect on binding of recombinant SARS-CoV-2 RBD protein Heparin Lyase treatment is included as control for HS dependent adhesion. **f-g,** effect of halofuginone treatment on the infection of SARS-CoV-2 spike protein pseudotyped virus in Hep3B (**f**) and Vero E6 (**g**) cells (n= 3-4 replicates per condition). **h**, Authentic SARS-CoV-2 virus infection of Hep3B cells treated with halofuginone. **i,** Infection of primary human bronchial epithelial cells, grown at an air-liquid interface, treated with halofuginone (Red and black colors represent the two different primary human bronchial epithelial cells isolates used). **j**, Relative cell viability of primary human bronchial epithelial cells. Data shown as mean ± S.D. Statistics performed by 1-way ANOVA and uncorrected Fisher’s LSD test (ns: *p* > 0.05, *: *p* ≤ 0.05, **: *p* ≤ 0.01, ***: *p* ≤ 0.001).

### Halofuginone Inhibits Heparan Sulfate Biosynthesis

The binding of SARS-CoV-2 spike protein to cells is HS-dependent^3^. HS is a linear polysaccharide attached to serine residues in HS proteoglycans (HSPGs)^21^. The HS polysaccharides consist of alternating residues of *N*-acetylated or *N*-sulfated glucosamine (GlcNAc or GlcNS) and either glucuronic acid (GlcA) or iduronic acid (IdoA) (Fig. 2a). Hence, we tested if halofuginone reduces spike protein binding by reducing HS presentation at the cell surface. Hep3B and Vero E6 cells were treated for 18 h with halofuginone and the effects on cellular HS were evaluated using the monoclonal antibody (mAb) (10E4) that recognizes a common epitope in HS (Fig. 2b & **Extended data Fig. 3**). Halofuginone dose-dependently reduced 10E4 binding in Hep3B (Fig. 2c), whereas 10E4 binding increased in Vero E6 upon halofuginone treatment (Fig. 2d). This directly correlates with the level of spike RBD binding and S protein pseudotyped virus infection in these cells (Fig. 1b-c). This finding suggests that halofuginone inhibits spike protein binding and SARS-CoV-2 viral attachment by altering cell surface HS content. Analysis of Hep3B cells showed that 0.5 µM halofuginone reduced cell surface HS in Hep3B cells by ∼4-fold (Fig. 2e). No difference was seen in the disaccharide composition following halofuginone treatment, suggesting that halofuginone affects total HS synthesis but does not alter HS specific sulfation (Fig. 2f). To test if the observed decrease in total HS was due to a decrease in availability of HSPG core proteins to carry HS, we lysed treated Hep3B cells, ran the lysates on SDS-PAGE and stained using an mAb (3G10) that recognizes a neo-epitope remaining on the proteoglycans (Fig. 2b,g) ^22^. The analysis revealed that halofuginone reduced the expression of core HSPGs in a dose-dependent manner (Fig. 2g & **Extended data Fig. 4**). These data suggest that halofuginone inhibits HS-mediated binding of the SARS-CoV-2 spike protein to cells by inhibiting the expression of HSPGs.

**Figure 2.**
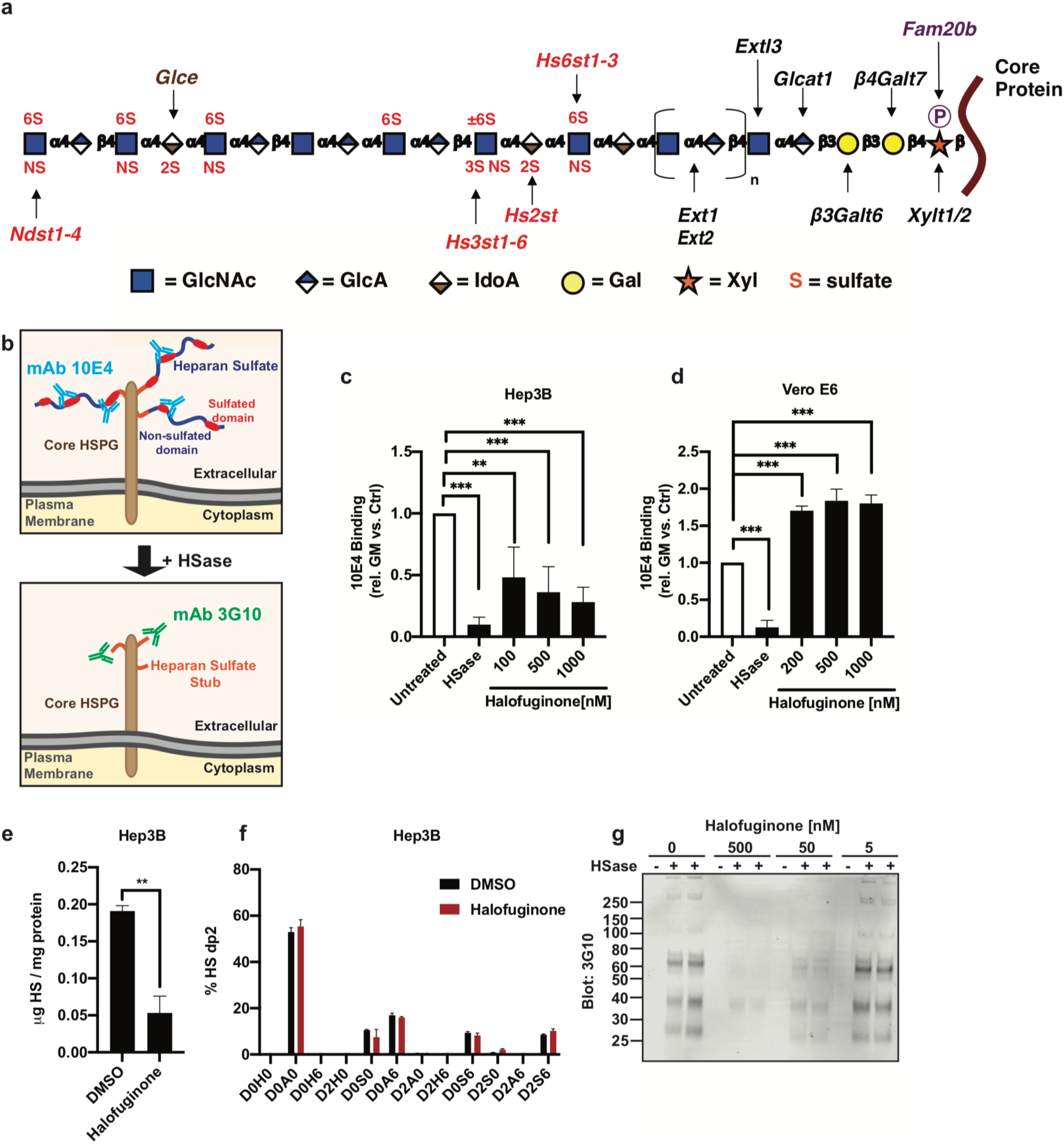
Halofuginone Inhibits Heparan Sulfate Biosynthesis. **a,** Schematic representation of HS biosynthesis. Genes required for priming and elongation are highlighted in black. Sulfotransferases and other modifying enzymes are highlighted in red. **b,** Schematic example of the interaction with the anti-HS 10E4 and 3G10 mAb’s. 10E4 recognizes sulfated HS polymer, while 3G10 recognizes the number of HS attachments sites by interacting with the stub left after heparin lyase (HSase) treatment. **c-d,** titration of halofuginone on Hep3B cells (**c**), Vero E6 (**d**), and its effect on cellular staining with anti-HS 10E4 mAb (n = 3 per group). **e,** quantitative determination of HS content in Hep3B cells treated with halofuginone. Absolute HS content was determined by disaccharide analysis using LC-MS (n = 3 per group). **f,** Qualitative distribution of specific sulfation patterns in Hep3B cells treated with halofuginone, seen in the LC-MS analysis (n = 3 per group). **g,** Quantification of functional HS binding sites in Hep3B cells treated with halofuginone. Binding sites are quantified in SDS-PAGE using mAb 3G10 that recognizes the linker tetrasaccharide bound to the HSPG core protein after hep lyase treatment. Data shown as mean ± S.D. Statistics performed by unpaired t test or 1-way ANOVA and uncorrected Fisher’s LSD test (ns: *p* > 0.05, *: *p* ≤ 0.05, **: *p* ≤ 0.01, ***: *p* ≤ 0.001, ****: *p* ≤ 0.0001).

To broadly examine the effects of halofuginone treatment, Hep3B cells were treated with vehicle or halofuginone at 200 or 500 nM for 6 h and 18 h and processed for RNA-sequencing (RNA-Seq). Halofuginone profoundly impacted the transcriptome (∼2700 differentially expressed genes), which increased with longer incubation time (**Extended data Fig. 5a-b**). Principal component analysis clearly segregated the response with respect to the duration of treatment (**Extended data Fig. 5a**). Analysis of halofuginone upregulated genes did not identify common pathways associated with antiviral or inflammatory responses (**Extended data Fig. 5c-d**)^23^. Additionally, no significant difference in expression of host factors exploited by SARS-CoV-2, such as *TMPRSS2* and *ACE2*, were observed (**Extended data Fig. 5f-g**). Gene annotation analysis of downregulated genes at 6 h showed that 200 nM halofuginone altered the expression of genes involved in glycoprotein biosynthesis and proteoglycan metabolic processes (**Extended data Fig. 5e**)^23^. A select group of core HSPGs were downregulated at the mRNA level (**Extended data Fig. 5f-g**). In particular, HSPGs *GPC2* and *SDC1* were significantly downregulated in conjunction with the HS biosynthetic enzymes *B3GAT3* and *EXTL3* (**Extended data Fig. 5f-g & 6**). Taken together, the data demonstrates that halofuginone suppresses the expression of proteins involved in HS and HSPG production.

### Halofuginone Inhibition of Spike Protein Binding to Heparan Sulfate is Independent of the Canonical Integrated Stress Response

Aminoacyl-tRNA-synthetases (AARS) catalyze the ATP-dependent synthesis of amino-acylated tRNAs via a two-step reaction involving an aminoacyl-adenylate intermediate with subsequent transfer to the cognate tRNAs^24^. Halofuginone and chemically unrelated compounds like prolyl-sulfamoyl adenosine (ProSA), specifically and potently inhibit human prolyl-tRNA synthetase (PRS) by blocking distinct portions of their ligand binding pockets (Fig. 3a-c)^15,25^. PRS inhibition can specifically suppress translation of proteins, such as collagens, that are enriched in prolines while having minimal effects on general protein synthesis^15^. However, PRS inhibition can also lead to GCN2-mediated activation of the Integrated Stress Response (ISR). GCN2 (gene symbol *EIF2AK4*) senses uncharged tRNAs and phosphorylates the eukaryotic transcription initiation factor 2*α* (eIF2*α*), leading to a general reduction in 5’cap-mediated RNA translation and selective translation of the eukaryotic transcription factor ATF4 and its target genes (Fig. 3a)^26^. These genes contain structural features in their 5’UTR allowing for selective translation in the presence of Ser51 phosphorylated eIF2*α*. The complexity and transient nature of the transcriptional and translational changes mediated by different eIF2*α* kinases of the ISR allows a cell to adapt and resolve various stress situations including amino acid starvation or unfolded protein stress^26^.

**Figure 3.**
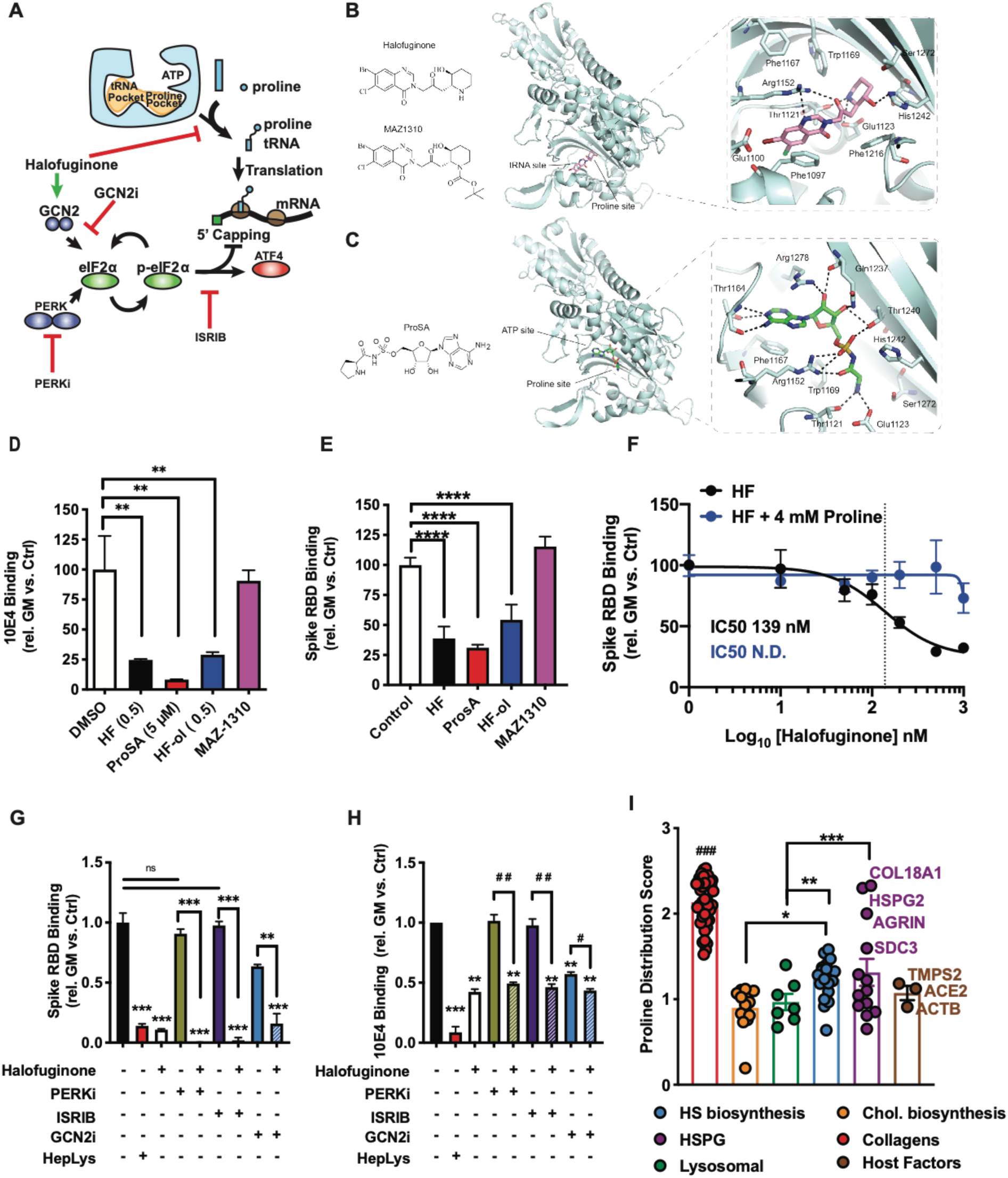
Halofuginone Inhibition of heparan sulfate presentation and Spike protein binding is not dependent of the integrated stress response. **a,** Chemical structure of halofuginone and negative control compound MAZ1310. Figure shows the interaction of halofuginone with the human prolyl-tRNA synthetase (PRS) active site, as resolved by X-ray crystallography (PDB: 4K88)^48^. **b,** Chemical structure of ProSA. Graphic shows the interaction of ProSA with human PRS (PDB: 5V58)^49^. **c,** Schematic representation of the mediators of the integrated stress response (ISR). **c-d,** Treatment of Hep3B cells with modulators of the PRS pathway at 0.5 µM (ProSA at 5 µM) and its effect on (**c**) HS presentation as measured by anti-HS mAb 10E4 binding and (**d**) spike RBD binding (**e**) by flow cytometry. Binding is represented as relative to non-treated control. **f,** Treatment of Hep3B cells with modulators of the halofuginone with or without 4mM proline and its effect on spike RBD binding by flow cytometry. Binding is represented as relative to non-treated control. **g,** Treatment of Hep3B cells with modulators of the ISR and its effect on HS presentation as measured by anti-HS mAb 10E4 binding in flow cytometry. **h,** Treatment of Hep3B cells with modulators of the ISR and its effect on SARS-CoV-2 recombinant RBD protein binding in flow cytometry. Binding is represented as relative to non-treated control. **i,** The distribution of proline distribution and density depicted as a proline distribution score for collagens, cholesterol biosynthetic proteins, lysosomal proteins, heparan sulfate (HS) biosynthetic proteins, heparan sulfate proteoglycans (HSPG) and viral host factor proteins (^##^*p* ≤ 0.01 collagens vs. all other protein classes). Data shown as mean ± S.D. Statistics performed by 1-way ANOVA and uncorrected Fisher’s LSD test (ns: *p* > 0.05, *: *p* ≤ 0.05, **: *p* ≤ 0.01, ***: *p* ≤ 0.001, ****: *p* ≤ 0.0001; ^#^:*p* ≤ 0.05, ^##^: *p* ≤ 0.01).

Interestingly, halofuginone induced a general ATF4-mediated ISR but did not activate the unfolded protein-induced ER stress response, as reported previously (**Extended data Fig. 5d**)^27^. To date, alterations in HS biosynthesis have not been identified as a hallmark of the ISR. To better understand the relationship between PRS inhibition and HS expression we tested related PRS inhibitor analogs to deconstruct the PRS pathway in relation to HS biosynthesis and spike RBD binding. Treatment with the non-cleavable and highly selective prolyl-AMP substrate analog ProSA at 5 μM prevented HS presentation and spike RBD binding to similar levels as treatment with halofuginone (500 nM), as illustrated by mAb 10E4 stain (Fig. 3c-d)^28^. Additionally, halofuginol (HFol), a halofuginone derivative that inhibits PRS, significantly decreased 10E4 and spike RBD binding while the MAZ1310 negative control compound, had no effect (Fig. 3d-e)^15,25^. Collectively, the data demonstrates that PRS inhibition is sufficient to reduce HS biosynthesis and spike RBD binding (Fig. 3d-e). Halofuginone competes with proline for the PRS active site^15^. Addition of excess proline to the media (4mM) of Hep3B cells prevented the ability of halofuginone to inhibit RBD protein binding to the cells (Fig. 3f).

To determine if halofuginone suppresses HS biosynthesis by activating the ISR we co-incubated halofuginone in the presence of a general inhibitor of the ISR, ISRIB (Integrated Stress Response inhibitor), selective eIF2*α* kinase inhibitors GCN2-IN-1 (GCN2i) targeting GCN2 (general control nonderepressible 2), or GSK2606414 (PERKi) targeting eIF2*α* kinase 3 (eIF2AK3), also known as protein kinase R-like endoplasmic reticulum kinase (PERK) (Fig. 3A)^26,29^. Neither PERKi nor ISRIB affected 10E4 or RBD binding, and neither had an effect on halofuginone inhibition (Fig. 3g-h). In contrast, GCN2i reduced 10E4 and RBD binding almost to the same extent as halofuginone activity (Fig. 3g-h), suggesting a role for GCN2 in the regulation of HS biosynthesis independent of the ISR. Neither GCN2-IN-1, GSK2606414, nor ISRIB reversed the halofuginone induced reduction in 10E4 or spike protein binding, suggesting that halofuginone does not suppress HS biosynthesis and spike binding by activating the ISR (Fig. 3g-h).

PRS inhibitors can selectively modulate the translational efficiency of proline-rich proteins, such as collagens (**Extended data Fig. 7**). Interestingly, HSPGs, such as agrin, perlecan, collagen 18 and syndecans 1 and 3, are relatively proline-rich compared to other proteins, such as TMPRSS2, ACE2 and lysosomal or cholesterol biosynthetic proteins (Fig. 3i and **Extended data Fig. 7-9; Extended data Table 1**). These observations suggest that inhibition of prolyl-tRNA charging could impair production of key membrane and extracellular matrix HSPGs (Fig. 3i and 2g). Together, these data suggest that PRS inhibitors, such as halofuginone, inhibit HS and proteoglycan expression both at the translational and transcriptional level.

### Halofuginone Inhibits infection and Replication by Authentic SARS-CoV-2

To determine if halofuginone inhibition of SARS-CoV-2 virus production was due to reduced viral entry or subsequent intra-host replication, Huh 7.5 cells were treated with 100 nM halofuginone or vehicle, either before, after, or before and after infection with SARS-CoV-2. Similarly, as in Hep3B halofuginone prevented productive infection of Huh 7.5 without affecting cell viability (Fig. 4a-b, and **Extended data Fig. 10)**. Pretreatment with halofuginone alone significantly reduced SARS-CoV-2 infection by ∼30-fold (Fig. 4a). However, halofuginone added after viral infection reduced the amount of secreted infectious virions by nearly 1000-fold (Fig. 4a). Intracellular viral RNA did not change when the cells were only treated before infection as expected, but viral RNA levels dramatically decreased 10- to 100-fold when halofuginone was present after infection (Fig. 4b). These observations suggest that halofuginone inhibits SARS-CoV-2 viral replication in addition to suppressing HS-dependent infection.

**Figure 4.**
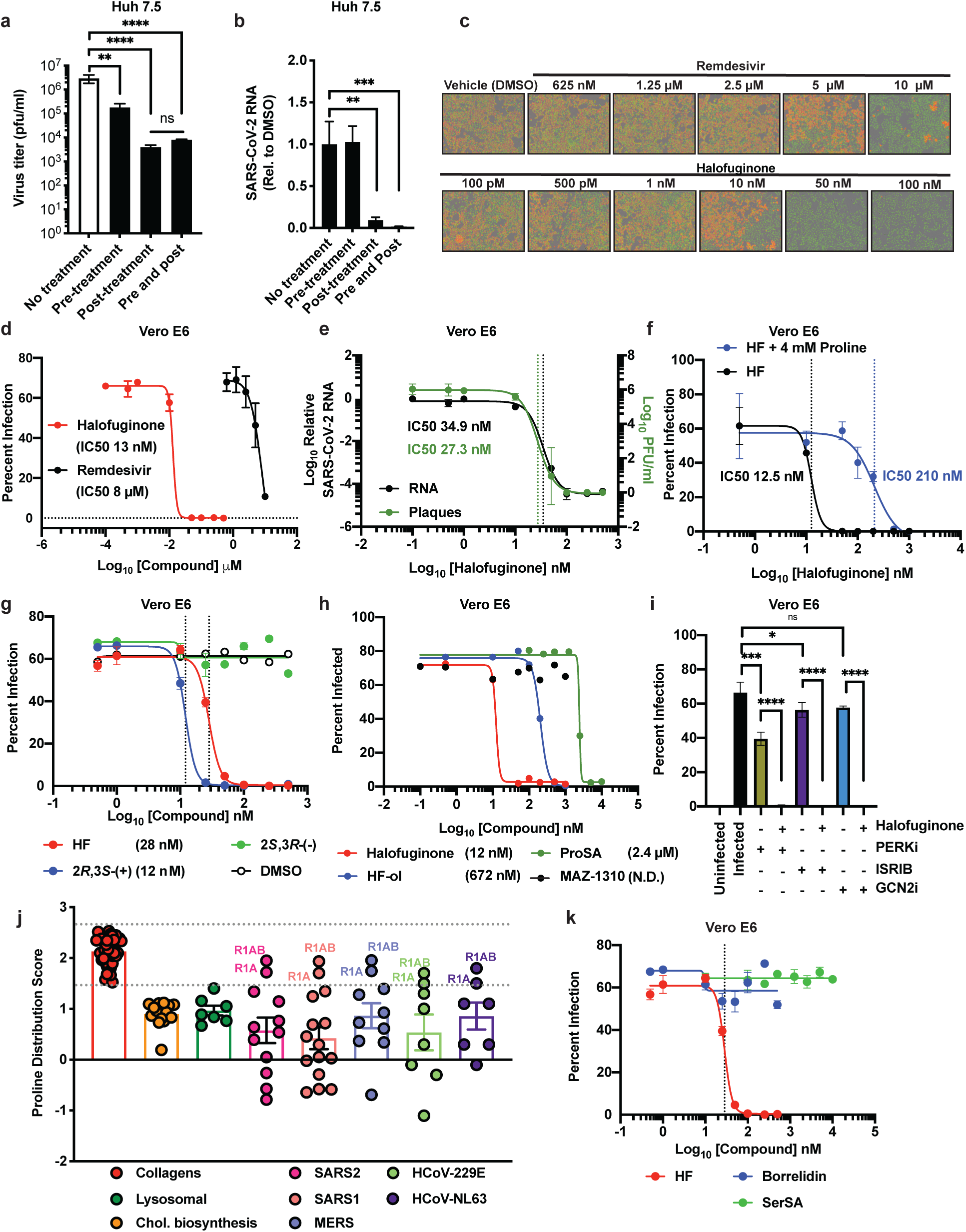
Halofuginone Inhibits infection and replication of authentic SARS-CoV-2. **a-b,** Authentic SARS-CoV-2 virus infection of Huh7.5 cells treated with Halofuginone. Huh7.5 cells treated with halofuginone pre- or post-infection, or both pre- and post-infection with authentic SARS-CoV-2 virus. Viral titers (**a**) and quantification of viral RNA (**b**) in the infected cells is shown. **c,** Immunofluorescent quantification of viral nucleocapsid (red) protein in Vero E6 cells treated with Halofuginone and Remdesivir and infected with authentic SARS-CoV-2 virus (nuclei = green). **d,** Authentic SARS-CoV-2 virus infection of Vero E6 cells treated with Halofuginone and Remdesivir as measured in flow cytometry. **e,** Quantification of plaque formation and viral RNA in Vero E6 cells treated with Halofuginone and Remdesivir and infected with authentic SARS-CoV-2 virus. **f,** Rescue experiment of the effect of halofuginone treatment on SARS-CoV-2 infection using excess proline. **g**, Treatment of Vero E6 cells with halofuginone and enantiomers and their effect on SARS-CoV-2 infection. **h,** Treatment of Vero E6 cells with modulators of the PRS pathway and their effect on SARS-CoV-2 infection. **i,** Treatment of Vero E6 cells with modulators of the ISR pathway and their effect on SARS-CoV-2 infection. **j,** The distribution of proline distribution and density depicted as a proline distribution score for collagens, cholesterol biosynthetic proteins, lysosomal proteins and viral SARS-CoV-2 (SARS2), SARS-CoV-1 (SARS1), MERS-CoV (MERS), HCoV-229E and HCoV-LN63 proteins(^##^*p* ≤ 0.01 collagens vs. all other protein classes). **k,** Treatment of Vero E6 cells with AARS inhibitors and their effect on SARS-CoV-2 infection. Data shown as mean ± S.D. Statistics performed by 1-way ANOVA and uncorrected Fisher’s LSD test (ns: *p* > 0.05, *: *p* ≤ 0.05, **: *p* ≤ 0.01, ***: *p* ≤ 0.001, ****: *p* ≤ 0.0001, ^##^: *p* ≤ 0.01.

Halofuginone did not decrease cellular HS or the binding of recombinant RBD protein in Vero E6 cells. However, given the effects of halofuginone treatment on viral replication we examined the effect of halofuginone on infection by authentic SARS-CoV-2 in Vero E6 cells as well. Halofuginone completely inhibited SARS-CoV-2 infectivity at 50 nM with an IC_50_ of 13 nM as measured by immunofluorescent (IF) detection of the nucleocapsid protein and plaque assays (Fig. 4c-d & **Extended data Fig. 11**). In comparison, Remdesivir had a calculated IC_50_ of 8 µM. Thus, halofuginone showed a ∼1,000-fold more potent inhibition of infection as compared to Remdesivir in this experimental setup (Fig. 4c-d). Similarly, halofuginone was 100-fold more potent compared to chloroquine (IC_50_ 1.9 µM) (**Extended data Fig. 12**). Moreover, halofuginone treatment reduced SARS-CoV-2 spike intracellular mRNA levels more than 20,000-fold with an IC_50_ of 34.9 nM (Fig. 4e).

We next examined if the impact of halofuginone on viral replication was dependent on PRS inhibition. Halofuginone competes with proline for the PRS active site^15^. Addition of excess proline to the media (4mM) of SARS-CoV-2 infected Vero E6 cells increased the IC_50_ of halofuginone from 12.5 nM to 210 nM (Fig 4f). Commercial halofuginone is a racemate of two, 2*S*,3*R*-(-) and 2*R*,3*S*-(+) enantiomers^30^. Hence, we evaluated if viral inhibition was due to on-target effects of the PRS targeting 2*R*,3*S*-(+) enantiomer (Fig. 4g)^30^. The 2*R*,3*S*-(+) inhibited SARS-CoV-2 infectivity with an IC_50_ of 12 nM compare to an IC_50_ of 28 nM for the racemic mixture. No inhibition was observed when using the non-targeting 2*S*,3*R*-(-) (Fig. 4g). Analogous, only the active 2*R*,3*S*-(+) enantiomer inhibited HS production and authentic SARS-CoV-2 spike RBD protein binding (**Extended data Fig. 13-14**).

Like halofuginone, other PRS inhibitors ProSA and halofuginol inhibited authentic SARS-CoV-2 infection of Vero E6 cells, although at significantly higher doses (Fig. 4h). The negative control compound, MAZ1310, did not (Fig. 4h). These results demonstrate that inhibition of PRS activity suppresses SARS-CoV-2 infection (Fig. 4h). Inhibitors of the ISR, as well as inhibitors of eIF2-alpha kinases PERK and GCN2 were unable to attenuate the halofuginone-mediated inhibition of SARS-CoV-2 infectivity (Fig. 4i). Hence, the observed antiviral effect is not dependent on ISR activation, consistent with the effects seen on the binding of recombinant RBD spike protein and mAb 10E4 binding (Fig. 3c-d).^31,32^ Upon cell entry, SARS-CoV-2 genomic RNA is translated into two large polyproteins, pp1a (>400 kDa) and pp1ab (>700 kDa) that undergo proteolytic processing into 11 or 16 non-structural proteins, respectively, many of which are required for RNA replication. Both pp1a (R1A) and pp1ab (R1AB), and to a lesser degree spike protein, have high proline contents comparable to collagens (Fig. 4j & **Extended data Fig. 15; Extended data table 1**). To probe the importance of the viral protein proline content versus general mRNA translation inhibition we evaluated two other AARS inhibitors, borrelidin and seryl-sulfamoyl adenosine (SerSA) that target the threonyl- and seryl-tRNA-synthetase, respectively. Niether of these AARS inhibitors were able to significantly attenuate viral replication (Fig. 4k) or Spike protein binding (**Extended data Fig. 16**), suggesting that proline translation is the Achilles heel for SARS-CoV-2 replication and possible other RNA viruses^33,34^.

## DISCUSSION

We show that halofuginone is a potent inhibitor of SARS-CoV-2 infection in numerous cell types, including primary human bronchial epithelial cells. This small molecule decreases HS-dependent spike protein binding, SARS-CoV-2 pseudovirus infection, and infection by authentic SARS-CoV-2. Interestingly, it also inhibits authentic SARS-CoV-2 infection post-entry by an HS-independent mechanism. Mechanistically, halofuginone suppresses SARS-CoV-2 infection by inhibiting the PRS, which could suppress the translation of long proline-rich host attachment factors, particularly HSPGs, and SARS-CoV-2 polyproteins pp1a and pp1ab that encode proteins required for viral replication. Thus, halofuginone is a potent host-targeting antiviral with dual inhibitory activity against SARS-CoV-2.

There is a desperate need for a potent, orally bioavailable antiviral for the treatment of COVID-19 that can be administered early in the disease to prevent hospitalization and the development of severe pulmonary disease. Here, we demonstrate that halofuginone is a potent inhibitor of SARS-CoV-2 infection with IC50 values in the low nanomolar range, in multiple *in vitro* models of infection. Halofuginone is orally bioavailable and reached an average C_max_ of 0.54 ng/ml (∼1.3 nM) or 3.09 ng/ml (∼7.4 nM) after a single administration of 0.5 mg or 3.5 mg doses in a phase I clinical trial^35^. The long half-life (∼30 h) leads to accumulation of halofuginone, with two- to three-fold higher exposure by day 15 of dosing^35^. Halofuginone widely distributed in tissues after administration in mice with the highest tissue concentrations in the lung and kidney^36^. Expressed as area under the curve, halofuginone exposure was more than 87-fold higher in the lung compared to plasma after a single intravenous injection in mice^36^. This suggests that although it may be difficult to obtain halofuginone plasma levels significantly above the IC_50_ values determined in this study, doses tested in phase I trials may be sufficient to achieve significant anti-SARS-CoV-2 activity in the lung and other organs infected by SARS-CoV-2^37–40^.

Excessive inflammation can contribute to inflammatory organ injury during severe COVID-19. There are multiple anti-inflammatory agents under evaluation for the treatment of COVID-19 and low dose dexamethasone recently demonstrated lower 28-day mortality in individuals receiving respiratory support^41^. In addition to the antiviral activity against SARS-CoV-2 that is described here, halofuginone has anti-inflammatory and anti-fibrotic activity that may provide yet another additive benefit to individuals with COVID-19 pneumonia^1,2,15,42–44^.

We demonstrate that other PRS inhibitors, ProSA and halofuginol, also inhibit infection by authentic SARS-CoV-2. Our data suggests that PRS inhibition may be particularly effective at suppressing the production of SARS-CoV-2 polyproteins pp1a and pp1ab that are long and proline-rich. Other positive-sense ssRNA viruses produce long polyproteins that may be similarly sensitive to PRS or other AARS inhibitors. Consistent with this idea, halofuginone demonstrates antiviral activity against future deleterious coronaviruses and other positive ssRNA viruses, including Chikungunya virus, Dengue virus, Zika virus, and Sindbis virus (**Extended data Fig. 17; Extended data table 1**)^45^.

In addition to halofuginone, other PRS inhibitors are being evaluated in humans, including DWN12088 that is in phase I clinical trials in Australia and anticipated to be used for the treatment of interstitial pulmonary fibrosis. Thus, evaluation of PRS inhibitors, halofuginone analogs, and other AARS inhibitors may lead to the identification of additional broad-spectrum antiviral agents. Viruses require host cell resources for replication and thus can be inhibited by therapeutics that target essential host factors. This approach could provide broad activity against diverse viruses while decreasing the risk of emerging viral resistance as the therapy is not directed against a specific virally encoded product. On the other hand, targeting host factors can lead to cytostatic and cytotoxicity, and we did observe the reported anti-proliferative effect of halofuginone in our assays (**Extended data Fig. 17**)^2^. Although inhibition of global translation could lead to excessive toxicity, inhibiting AARSs could provide a more precise way of inhibiting viral proteins and thus limit toxicity. It appears that normal cells are relatively tolerant of decreased AARS levels with minimal effects on global translation. Individuals who are heterozygous carriers for recessive inactivating AARS mutations linked to hypomyelinating leukodystrophy do not display disease phenotypes^46^.

In conclusion, we identified halofuginone as an antiviral with potent inhibitory activity against SARS-CoV-2 infection in multiple human cell types. We show that halofuginone reduces both HS and HSPG biosynthesis, that are required for viral adhesion. In addition, we find that the inhibitory capacity of halofuginone on SARS-CoV-2 infection is not limited to HS reduction, but also shows potent inhibition in the viral replication stage. Mechanistically, our data suggests that these effects are caused by PRS inhibition and its effect on production of proline-rich proteins and not dependent on ISR activation. This observation is in agreement with previous studies reporting that coronaviruses overcome the inhibitory effects of eIF2*α* phosphorylation on viral mRNA translation ^31,32^. Furthermore, many viruses, including coronaviruses (SARS1 and SARS2), shutdown host translation using nonstructural protein 1 (Nsp1) and activate eIF2*α* associated PERK dependent stress responses that benefit the virus^47^. Halofuginone may prevent SARS-CoV-2 from circumventing these mechanisms of host protein translational shutdown.

Halofuginone oral administration has been evaluated in a phase I clinical trial in humans and based on pharmacological studies in mice is distributed to SARS-CoV-2 target organs, including the lung. In conclusion, we show that halofuginone is a potent inhibitor of SARS-CoV-2 infection which emphasizes its potential as an effective treatment for COVID-19 in the clinic. Beyond its antiviral activity, halofuginone has potent anti-inflammatory and anti-fibrotic properties that may provide additive benefit in the treatment of COVID-19 pneumonia. Based on this in vitro preclinical data we believe halofuginone could be an effective antiviral and antifibrotic agent for the treatment of individuals with COVID-19.

## ACKNOWLEDGMENTS

We thank John Christianson, University of Oxford, and Matthew Frieman at the University of Maryland for discussions and helpful comments, and Alexandra Efstrathiou, University of Oxford, for technical assistance.

## Author contributions

P.L.S.M.G., A.F.C., D.R.S., T.M.C. and C.N. conceived the project; P.L.S.M.G., A.F.C., J.D.E., J.J., S.T., B.A.C., S.L., R.M., B.A.C., K.Z., U.O., D.R.S., T.M.C. R.J. W. and C.N designed and performed the experimental work.; A.D., A.P.C., A.E.C., A.F.G, J.K.C.K., A.N., S.A.M., G.J.G., C.B.S, C.J.N., E.R.G., J.E.D, C.J.N, E.R.G, E.M.K., R.N.M. and A.P. performed the experimental work.: X.Y., Y.P., J.F., B.H., T.M.C., S.K.C., M.A.T., N.C.P, A.G.S. J.A.M., T.A., C.B, M.D., R.M.E. and R.M. supplied reagents. P.L.S.M.G., A.F.C. D.R.S., T.M.C., J.K.C.K., J.D.E., U.O., R.M., J.A.M., M.D., R.M.E., A.P.C., M.P, C.J., B.A.C., K.V.G. and S.T. analyzed and discussed the data.; T.M.C., D.R.S., A.F.C. and P.L.S.M.G, wrote the manuscript. All authors discussed the experiments and results, read, and approved the manuscript.

## Materials & Correspondence

Correspondence should be addressed to: Philip L.S.M. Gordts, Department of Medicine, University of California, La Jolla, CA 92093, USA; +1(858)246-0994, Aaron F. Carlin, Department of Medicine, University of California, La Jolla, CA 92093, USA; +1(858)246-3261

## METHODS

### Reagents

SARS-CoV-2 (2019-nCoV) Spike Protein (RBD, His Tag) (Sino Biologicals, 40592-V08B), and proteins were biotinylated using EZ-Link™ Sulfo-NHS-Biotin, No-Weigh™ (Thermo Fisher Scientific, A39256). Heparin lyases were from IBEX pharmaceuticals, Heparinase I (IBEX, 60-012), Heparinase II (IBEX, 60-018), Heparinase III (IBEX, 60-020). Protein production reagents included, Pierce™ Protein Concentrators PES Thermo Scientific™, Pierce™ Protein Concentrator PES (Thermo Fisher Scientific, 88517) and Zeba™ Spin Desalting Columns, 40K MWCO, 0.5 mL (Thermo Fisher Scientific, 87766). ISR pathway inhibitors used are ISRIB (Sigma, SML0843), GSK2606414 (Sigma, 516535), and A-92 (Axon, 2720). Antibodies used were, anti-Spike antibody [1A9] (GeneTex, GTX632604), anti-Nucleocapsid antibody (GeneTex, GTX135357), Anti-HS (Clone F58-10E4) (Fisher Scientific, NC1183789), and 3G10 (BioLegend, 4355S). Secondary antibodies were, Anti-His-HRP (Genscript, A00612), Avidin-HRP (Biolegend, 339902), and Streptavadin-Cy5 (Thermo Fisher, SA1011). Luciferase activity was monitored by Bright-Glo^TM^ (Promega, E2610). All cell culture medias and PBS where from Gibco.

### Cell Culture

Huh-7.5 cells ^1^ were generously provided by Charles M. Rice (Rockefeller University, New York, NY). Hep3B, BHK-15 and Calu-3 cells were from ATCC and were grown in DMEM medium containing 10% % fetal bovine serum, and 100 IU/ml of penicillin, 100 µg/ml of streptomycin sulfate and nonessential amino acids. BHK-15 were grown in Modified Eagle’s supplemented with 10% fetal bovine serum and nonessential amino acids (Gibco, #11140-050). Hep3B, A375 and Vero E6 cells were from ATCC. The Hep3B cells carrying mutations in HS biosynthetic enzymes was derived from the parent ATCC Hep3B stock, and have been described previously^2^. All cells were supplemented with 10% FBS, 100 IU/ml of penicillin and 100 µg/ml of streptomycin sulfate and grown under an atmosphere of 5% CO2 and 95% air. Cells were passaged before 80% confluence was reached and seeded as explained for the individual assays. Cell viability was measured using LDH leakage (Promega) or alamarBleu (Thermo Fisher).

### SARS-CoV-2 spike RBD protein production

Recombinant SARS-CoV-2 RBD (GenBank: MN975262.1; amino acid residues 319-529) was cloned into pVRC vector containing a HRV 3C-cleavable C-terminal SBP-His_8X_ tag was produced in ExpiCHO cells by transfection of 6 x10^6^ cells/ml at 37 °C with 0.8 μg/ml of plasmid DNA using the ExpiCHO expression system transfection kit in ExpiCHO Expression Medium (ThermoFisher). One day later the cells were fed with ExpiCho Feed, treated with ExpCho Enhancer, and then incubated at 32 °C for 11 days. The conditioned medium was mixed with cOmplete EDTA-free Protease Inhibitor (Roche). Recombinant protein was purified by chromatography on a Ni^2+^ Sepharose 6 Fast Flow column (1 ml, GE LifeSciences). Samples were loaded in ExpiCHO Expression Medium supplemented with 30 mM imidazole, washed in a 20 mM Tris-Cl buffer (pH 7.4) containing 30 mM imidazole and 0.5 M NaCl. Recombinant protein was eluted with buffer containing 0.5 M NaCl and 0.3 M imidazole. The protein was further purified by size exclusion chromatography (HiLoad 16/60 Superdex 200, prep grade. GE LifeSciences) in 20 mM HEPES buffer (pH 7.4) containing 0.2 M NaCl.

### SARS-CoV-2 Spike RBD Biotinylation

For binding studies, recombinant SARS-CoV-2 Spike RBD protein (RBD, His Tag) was conjugated with EZ-Link^TM^ Sulfo-NHS-Biotin (1:3 molar ratio; Thermo Fisher) in Dulbecco’s PBS at room temperature for 30 min. Glycine (0.1 M) was added to quench the reaction and the buffer was exchanged for PBS using a Zeba spin column (Thermo Fisher).

### Flow cytometry

Cells at 50-70% confluence were treated with halofuginone or other inhibitors for 18-24 hrs. The cells were lifted with PBS containing 10 mM EDTA (Gibco) and washed in PBS containing 0.1% BSA. The cells were seeded into a 96-well plate at 10^5^ cells per well. As a control for HS-dependent spike protein adhesion, a portion of the cells were treated with HSase mix (2.5 mU/ml HSase I, 2.5 or 5 mU/ml HSase II, and 5 mU/ml HSase III; IBEX) for 30 min at 37 °C in PBS containing 0.1% BSA. Cells were stained for 30 min at 4°C with biotinylated spike RBD protein (20 µg/ml) or 10E4 mAB in PBS containing 0.1% BSA. The cells were washed twice and then reacted for 30 min at 4°C with Streptavadin-Cy5 (Thermo Fisher; 1:1000 dilution) or anti-mouse IgG-Alexa488, respectively, in PBS containing 0.5% BSA. The cells were washed twice and then analyzed using a FACSCalibur or a FACSCanto instrument (BD Bioscience). All experiments were done a minimum of three separate times in three technical replicates. Data analysis was performed using FlowJo software and statistical analyses were done in Prism 8 (GraphPad).

### HS and CS purification from cell surface of Hep3B cells

Fresh Hep3B cells were was washed in PBS, lifted with trypsin, and centrifuged for 5 min a 500 x g. The trypsin supernatants were saved for GAG analysis, and the cell pellets were washed in complete media, followed by a PBS wash, and saved for BCA analysis (Pierce™ BCA Protein Assay Kit, Themo Fisher), according to vendor specifications. The trypsin-released GAGs were treated with 1 mg/mL Pronase (Streptomyces griseus, Sigma Aldrich) and 0.1% Triton X-100 and 10 mM CaCl_2_, and incubated at 37°C, overnight with shaking. The samples were centrifuged at 20,000 x g for 20 min and the supernatant was mixed 1:10 with equilibration buffer (50 mM sodium acetate, 0.2 M NaCl, 0.1% Triton X-100, pH 6) and loaded onto a DEAE Sephacel column (GE healthcare) equilibrated with equilibration buffer. The column was washed with wash buffer (50 mM sodium acetate, 0.2 M NaCl, pH 6.0) and bound GAGs were eluted with elution buffer (50 mM sodium acetate, 2.0 M NaCl, pH 6.0). The eluate was desalted using PD-10 columns (GE Healthcare) and lyophilized until dry.. The GAG pellets were reconstituted in DNase buffer (50 mM Tris, 50 mM NaCl, 2.5 mM MgCl_2_, 0.5 mM CaCl_2_, pH 8.0) with 20 kU/mL DNase I (Deoxyribonuclease I from Bovine Pancreas, Sigma Aldrich) and incubated for 2 hr at 37°C, shaking. Finally, the GAGs were purified over a DEAE column and precipitated by PD-10 chromatography ^3^.

### HS and CS digestion and MS analysis

For HS quantification and disaccharide analysis, purified GAGs were digested with a mixture of heparin lyases I-III (2 mU each) for 2 hr at 37 °C in lyase buffer (40 mM ammonium acetate and 3.3 mM calcium acetate, pH 7.0). For CS quantification and disaccharide analysis, the GAGs were digested with 20 mU/mL Chondroitinase ABC (from Proteus vulgaris, Sigma Aldrich) and incubated for 2 hr at 37°C in lyase buffer (50 mM Tris and 50 mM NaCl, pH 8.0). The reactions were dried in a centrifugal evaporator and tagged by reductive amination with [^12^C_6_]aniline. The HS and CS samples were analyzed by liquid chromatography (LC) coupled to tandem mass spectrometry (MS/MS) and quantified by inclusion of [^13^C_6_]aniline-tagged standard HS disaccharides (Sigma-Aldrich), as described (Lawrence et al., 2008). The samples were separated on a reverse phase column (TARGA C18, 150 mm x 1.0 mm diameter, 5 µm beads, Higgins Analytical, Inc.) using 5 mM dibutylamine as an ion pairing agent (Sigma-Aldrich), and ions were monitored in negative mode. Separation was performed using the same gradient, capillary temperature, and spray voltage as described (Lawrence et al., 2008). The analysis was done on an LTQ Orbitrap Discovery electrospray ionization mass spectrometer (Thermo Scientific) equipped with an Ultimate 3000 quaternary HPLC pump (Dionex).

### Western Blot Analysis

Cells were lysed using RIPA buffer, and protein was quantified using a BCA assay. Protein was analyzed by SDS-PAGE on 4–12% Bis-Tris gradient gels (NuPage; Invitrogen) with an equal amount of protein loading. Proteins were visualized after transfer to Immobilon-FL PVDF membrane (Millipore). Membranes were blocked with Odyssey blocking buffer (LI-COR Biosciences) for 30 min and incubated overnight at 4°C with 3G10 antibodies. Mouse antibodies were incubated with secondary Odyssey IR dye antibodies (1:14,000) and visualized with an Odyssey IR imaging system (LI-COR Biosciences).

### RNA-seq library preparation

Cells were lysed in Trizol and total RNA was extracted using the Direct-zol kit (Zymo Research, CA USA). On column DNA digestion was also performed with DNAse treatment. Poly(A) RNA was selected using the NEBNext Poly(A) mRNA Magnetic Isolation module (New England Biolabs) and libraries were prepared using the NEBNext Ultra Directional RNA Library Prep Kit (New England Biolabs) and sequenced using a NextSeq 500 (Illumina). Samples were sequenced at a minimum depth of 15 million reads per sample, paired end with a read length of 2×41bp.

### RNA sequencing data analysis

A computational pipeline was written calling scripts from the CGAT toolkit to analyse the RNA sequencing data (https://github.com/cgat-developers/cgat-flow) ^4,5^. Briefly, FASTQ files were generated and assessed for quality using FASTQC, aligned to GRCh38 (hg38) and then aligned to the transcriptome using hisat2 v2.1.0 ^6^. To count mapped reads to individual genes, featurecounts v1.4.6, part of the subreads package ^7^, was used. Differential gene expression analysis was performed using DESeq2 using treatment and time as factors in the model. Genes were considered to be differentially expressed based on log2 fold change and p-value < 0.05. R scripts used to analyze the transcriptomic data are available through GitHub (https://github.com/Acribbs/deseq2_report). Motif enrichment was performed using homer as described before ^8,9^.

### Preparation and infection by pseudotyped VSV

Vesicular Stomatitis Virus (VSV) pseudotyped with spike proteins of SARS-CoV-2 were generated according to a published protocol ^10^. Briefly, HEK293T, transfected to express full length SARS-CoV-2 spike proteins, were inoculated with VSV-G pseudotyped ΔG-luciferase or GFP VSV (Kerafast, MA). After 2 hr at 37°C, the inoculum was removed and cells were refed with DMEM supplemented with 10% FBS, 50 U/mL penicillin, 50 µg/mL streptomycin, and VSV-G antibody (I1, mouse hybridoma supernatant from CRL-2700; ATCC). Pseudotyped particles were collected 20 hr post-inoculation, centrifuged at 1,320 × g to remove cell debris and stored at −80°C until use. Cells were seeded at 10,000 cells per well in a 96-well plate. The cells were then treated with Halufuginone for 16hrs. As a control for HS dependent infection some cells were treated with HSases for 30 min at 37 °C in serum-free DMEM. Culture supernatant containing pseudovirus (20-100 µL) was adjusted to a total volume of 100 µL with PBS, HSase mix or the indicated inhibitors and the solution was added to the cells. After 4 hr at 37 °C the media was changed to complete DMEM. The cells were then incubated for 16 hr to allow expression of the luciferase gene. Cells were analyzed for infection by Bright-Glo^TM^ (Promega) using the manufacturers protocol. Briefly, 100 µL of luciferin lysis solution was added to the cells and incubated for 5 min at room temperature. The solution was transferred to a black 96-well plate and luminescence was detected using an EnSpire multimodal plate reader (Perkin Elmer). Data analysis and statistical analysis was performed in Prism 8.

### SARS-CoV-2 infection

SARS-CoV-2 isolate USA-WA1/2020 (BEI Resources) was propagated and infectious units quantified by plaque assay using Vero E6 (ATCC) cells. Approximately 10e4 Vero E6 cells per well were seeded in a 96 well plate and incubated overnight. The following day, cells were washed with PBS and 100uL of SARS-CoV-2 (MOI 0.5) diluted in serum free DMEM was added per well and incubated 1 h at 37°C with rocking every 10-15 min. After 1 h, virus was removed, cells washed with PBS and compounds or controls were added at the indicated concentrations. In experiments using inhibitors of the ISR or its related kinases, these compounds were added after viral infection and incubated for 1 h prior to the addition of HF. For viral RNA quantification, cells were washed twice with PBS and lysed in 200ul TRIzol (ThermoFisher). For immunofluorescence, cells were washed twice with PBS and incubated in 4% formaldehyde for 30 minutes at room temperature. For plaque assays, supernatant was removed and stored at −80 until plaque assays were performed.

### Flavivirus and alphavirus infection

Sindbis virus (SINV) with mCherry-tagged NSP3 was propagated on BHK-15 cells ^11^. Uganda strain of Zika virus (ZIKV) MR 766 encoding the fluorescent protein Venus from the genome (a kind gift from Matthew Evans, Schwarz 2016) was propagated on Huh 7.5 cells ^12^. Huh 7.5 and BHK-15 monolayers were pretreated with various halofuginone concentrations for 16 hours and infected with SINV or ZIKV at MOI of 0.1 for two hours in the presence of the compound. Subsequently, media over cells were replaced with fresh media containing halofuginone and incubated for 12 hours for SINV and 24 hours for ZIKV. The cell culture supernatants were collected at 12 hours post infection for SINV and 24 hours post infection for ZIKV and the virus titers were determined by fluorescent focus assay (FFT).

### Methods for human bronchial epithelial cell ALI generation and infection

#### Air Liquid Interface

Human Bronchial Epithelial Cells (HBECs, Lonza) were cultured in T75 flasks in PneumaCult-Ex Plus Medium according to manufacturer instructions (StemCell Technologies). To generate air-liquid interface cultures, HBECs were plated on collagen I-coated 24 well transwell inserts with a 0.4-micron pore size (Costar, Corning) at 5×10^4^ cells/ml. Cells were maintained for 3-4 days in PneumaCult-Ex Plus Medium until confluence, then changed to PneumaCult-ALI Medium (StemCell Technologies) containing ROCK inhibitor (Y-27632, Tocris) for 4 days. Fresh medium, 100 µl in the apical chamber and 500 µl in the basal chamber, was added daily. At day 7, the medium in the apical chambers was removed, and the basal chambers were changed every 2-3 days with apical washes with PBS every week for 28 days.

#### Human bronchial epithelial cell ALI Infection

The apical side of the HBEC ALI culture was gently washed three times with 200 µl of phosphate buffered saline without divalent cations (PBS-/-). An MOI of 0.5 of SARS-CoV-2 live virus in 100 µl total volume of PBS was added to the apical chamber with either DMSO, Heparinase or or various halofuginone concentrations. Cells were incubated at *37◦C* and 5% CO2 for 4 hours. Unbound virus was removed, the apical surface was washed and the compounds were re-added to the apical chamber. Cells were incubated for another 20 hours at *37◦C* and 5% CO2. After inoculation, cells were washed once with PBS-/- and 100 µl TrypLE (ThermoFisher) was added to the apical chamber then incubated for 10 min in the incubator. Cells were gently pipetted up and down and transferred into a sterile 15 ml conical tube containing neutralizing medium of DMEM + 3% FBS. TrypLE was added again for 3 rounds of 10 minutes for a total of 30 min to clear transwell membrane. Cells were spun down and resuspended in PBS with Zombie UV viability dye for 15 min in room temp. Cells were washed once with FACS buffer then fixed in 4% PFA for 30 min at room temp. PFA was washed off and cells were resuspended in PBS.

### Virus plaque assays

Confluent monolayers of Vero E6 or Hep3B cells were infected with SARS-CoV-2 at an MOI of 0.1. After one hour of incubation at 37 °C, the virus was removed, and the medium was replaced. After 48 hr, cell culture supernatants were collected and stored at −80°C. Virus titers were determined by plaque assays on Vero E6 monolayers. In short, serial dilutions of virus stocks in Dulbecco’s Modified Essential Media (DMEM) (Corning, #10-014-CV) were added to Vero E6 monolayers on 12-well plates and incubated 1 hr at 37 °C with rocking every 10-15 min. The cells were subsequently overlaid with MEM containing 0.6% agarose (ThermoFisher Scientific, #16500-100), 4% FBS, non-essential amino acids, L glutamine, and sodium bicarbonate and the plates were incubated at 37 °C under an atmosphere of 5% CO_2_/95% air for 48 hr. The plates were fixed with a mixture of 10% formaldehyde and 2% methanol (v/v in PBS) for 24 hr. Agarose overlays were removed, and the monolayer was washed once with PBS and stained with 0.025% Crystal Violet prepared in 2% ethanol. After 15 min, Crystal Violet was removed, and plaques were counted to determine the virus titers. Plaque assays were performed and counted by a blinded experimenter. All work with SARS-CoV-2 was conducted in Biosafety Level-3 conditions either at the University of California San Diego or at the Eva J Pell Laboratory, The Pennsylvania State University, following the guidelines approved by the Institutional Biosafety Committees.

### RNA extraction, cDNA synthesis and qPCR

RNA was purified from TRIzol lysates using Direct-zol RNA Microprep kits (Zymo Research) according to manufacturer recommendations that included DNase treatment. RNA was converted to cDNA using the iScript cDNA synthesis kit (BioRad) and qPCR was performed using iTaq universal SYBR green supermix (BioRad) and an ABI 7300 real-time pcr system. cDNA was amplified using the following primers RPLP0 F – GTGTTCGACAATGGCAGCAT; RPLP0 R – GACACCCTCCAGGAAGCGA; SARS-CoV-2 Spike F – CCTACTAAATTAAATGATCTCTGCTTTACT; SARS-CoV-2 Spike R – CAAGCTATAACGCAGCCTGTA. Relative expression of SARS-CoV-2 Spike RNA was calculated by delta-delta-Ct by first normalizing to the housekeeping gene RPLP0 and then comparing to SARS-CoV-2 infected Vero E6 cells that were untreated (reference control). Curves were fit and inhibitory concentration (IC) IC50 and IC90 values calculated using Prism 8.

### Immunofluorescence imaging and analysis

Formaldehyde fixed cells were washed with PBS and permeabilized for immunofluorescence using BD Cytofix/Cytoperm according to the manufacturers protocol for fixed cells and stained for SARS-CoV-2 with a primary anti-Nucleocapsid antibody (GeneTex GTX135357) followed by a secondary Goat anti Rabbit AF594 antibody (ThermoFisher A-11037) and nuclei stained with Sytox Green. Five or eight images per well were obtained using an Incucyte S3 (Sartorius) or Nikon Ti2-E microscope equipped with a Qi-2 camera and Lumencor Spectra III light engine respectively. The percent infected cells were calculated using built-in image analysis tools for the Incucyte S3. For images acquired with the Nikon Ti2, images were analysed using the Fiji distribution of ImageJ (PMID 22743772) and the DeepLearning plugin StarDist (Schmidt et al, 2012 – see below) as follows. Channels were separated and the Sytox Green-stained nuclei were segmented using StarDist to generate individual masks for each nucleus. The AF594 channel (Nucleocapsid) was selected by removing background with a rolling ball of 50 pixels. A median filter (sigma=10) was applied to facilitate easier thresholding of AF594 signal. The resulting image was thresholded and a mask was generated representing all positive staining. Positive nuclei were selected by firstly eroding the StarDist generated nuclei mask by 2 pixels to reduce potential overlap with non-positive stain. Binary reconstruction was then carried out between the resulting mask and the AF594 mask. Uwe Schmidt, Martin Weigert, Coleman Broaddus, and Gene Myers. Cell Detection with Star-convex Polygons. International Conference on Medical Image Computing and Computer-Assisted Intervention (MICCAI), Granada, Spain, September 2018.

### Proline distribution analysis

Protein amino acid sequences were downloaded from UniProtKB ^13^. Proline distribution was analyzed using custom code in R (v3.6.0) ^14^, including commands from the packages dplyr ^15^ tidyr ^16^, and stringr ^17^. All plots were generated in ggplot2 ^18^. For individual protein plots, histograms were constructed with geom_histogram(binwidth = 1) and kernel density estimations (KDEs) were constructed with geom_density(kernel = “gaussian”) and added as custom annotations (ggpubr package ^19^). The bandwidth of KDEs for individual plots was assessed separately for each protein distribution using maximum likelihood cross-validation ^20,21^ with the h.mlcv command from the kedd package ^22^. The proline distribution score was calculated using the following formula:

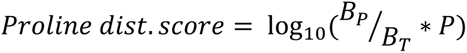

where B_P_ is the number of 10-amino acid blocks in a protein that contain one or more prolines (a protein with length 100 amino acids has 10 blocks); B_T_ is the total number of 10-amino acid blocks in a protein; and P is the total number of prolines in a protein. B_P_ values were obtained using custom code in R.

### Data and Code availability

Data are deposited in the National Centre for Biotechnology Information GEO database under the accession number (GSE157036). Depicted structures have been solved and deposited: halofuginone PDB ID 4K88, ProSA PDB ID 5V58, and NCP22 PDB ID 5VAD. The custom-scripted macro used for automated image analysis of live cell imaging data is available from https://doi.org/10.26180/5f508284eb365. Code and files used for proline analysis can be found on https://github.com/jkccoker/Proline_analysis.

---

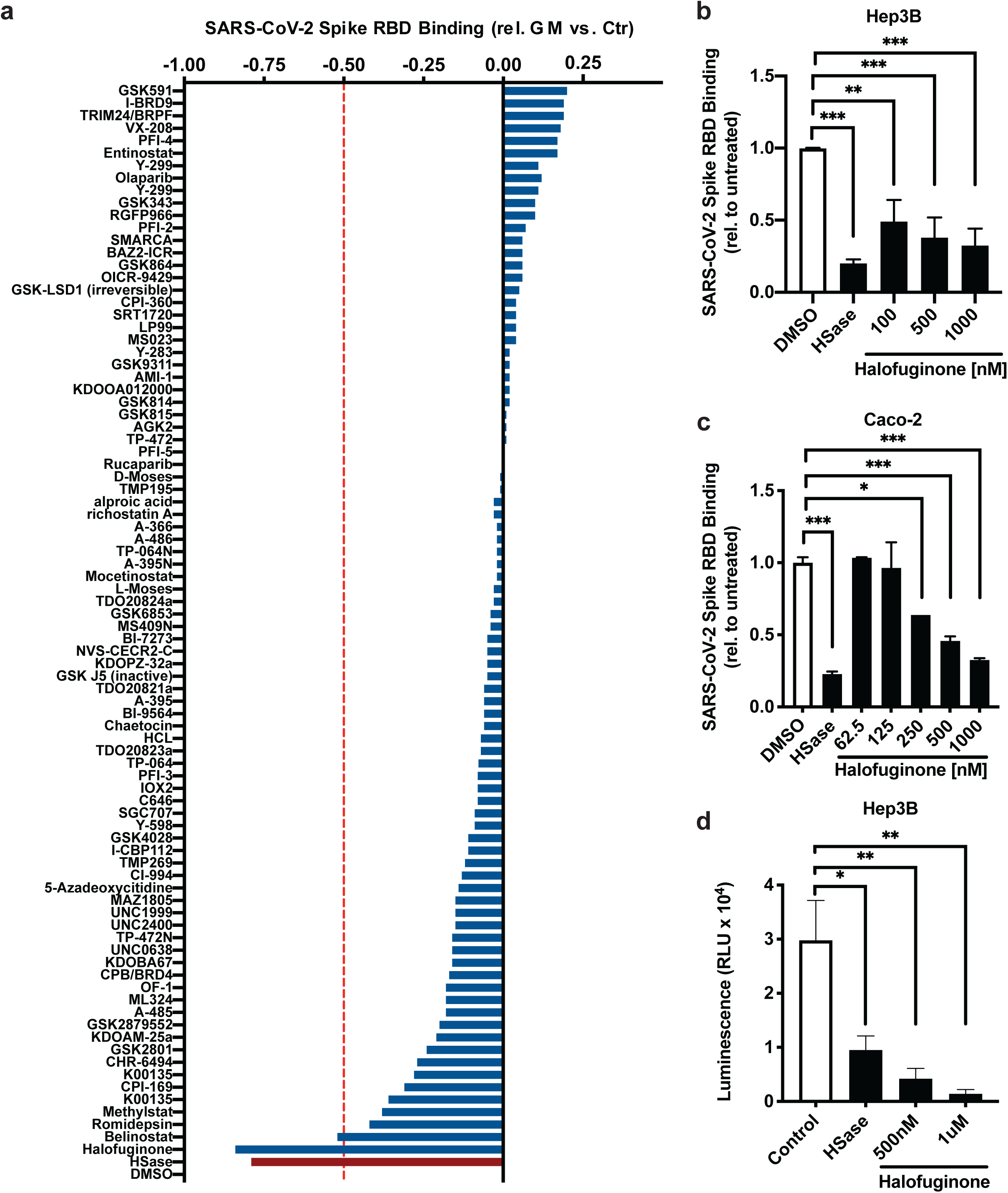

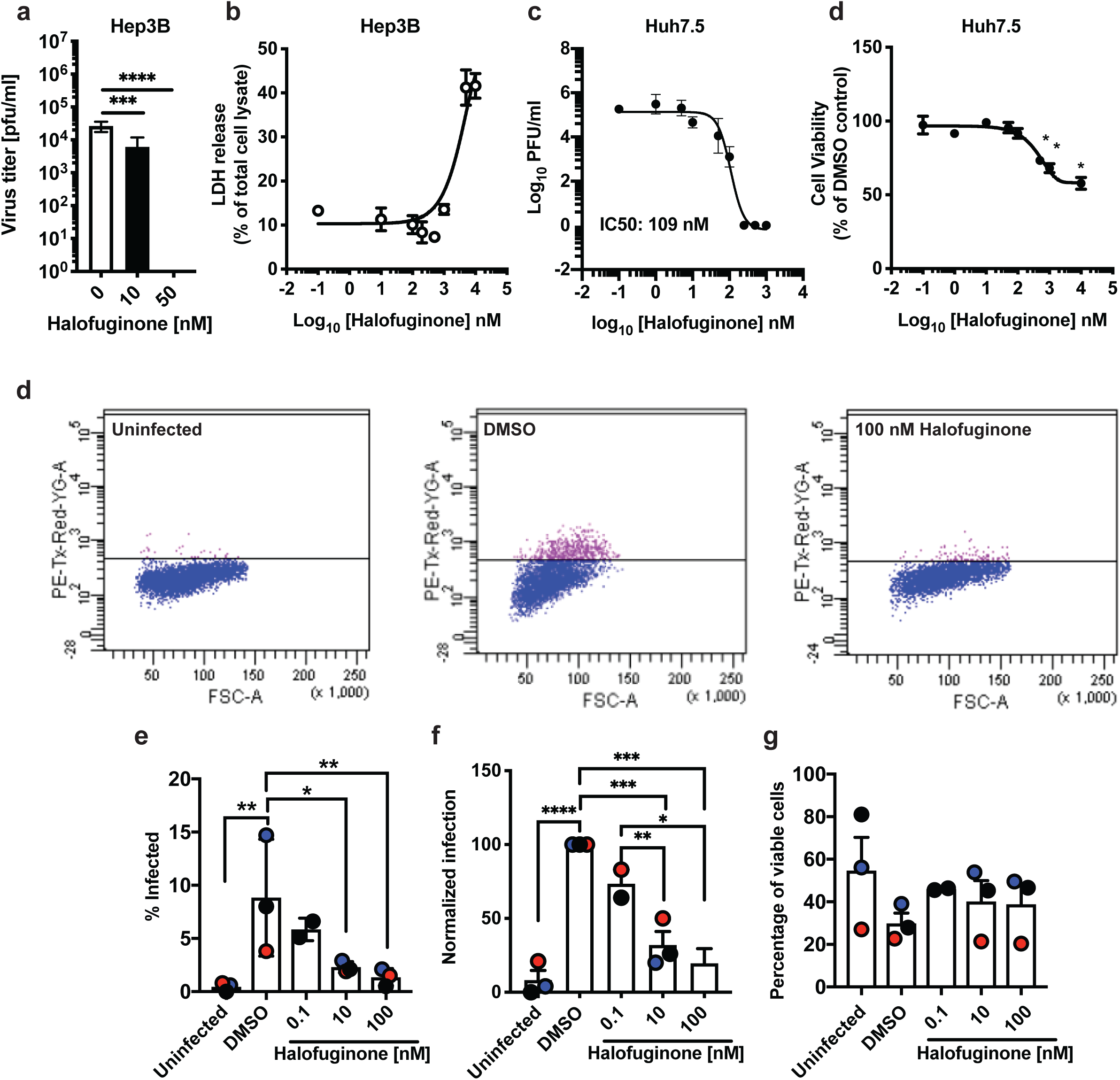

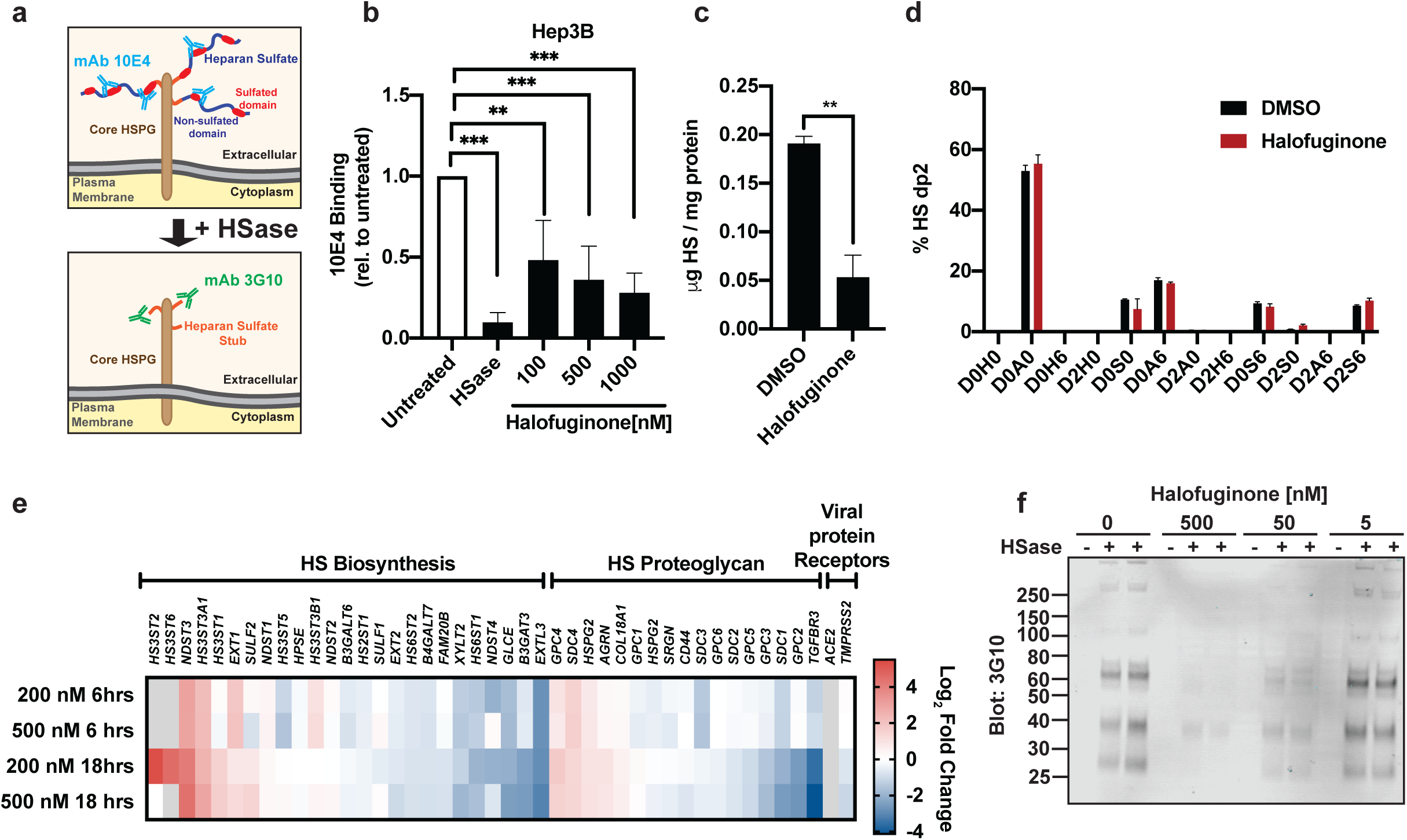

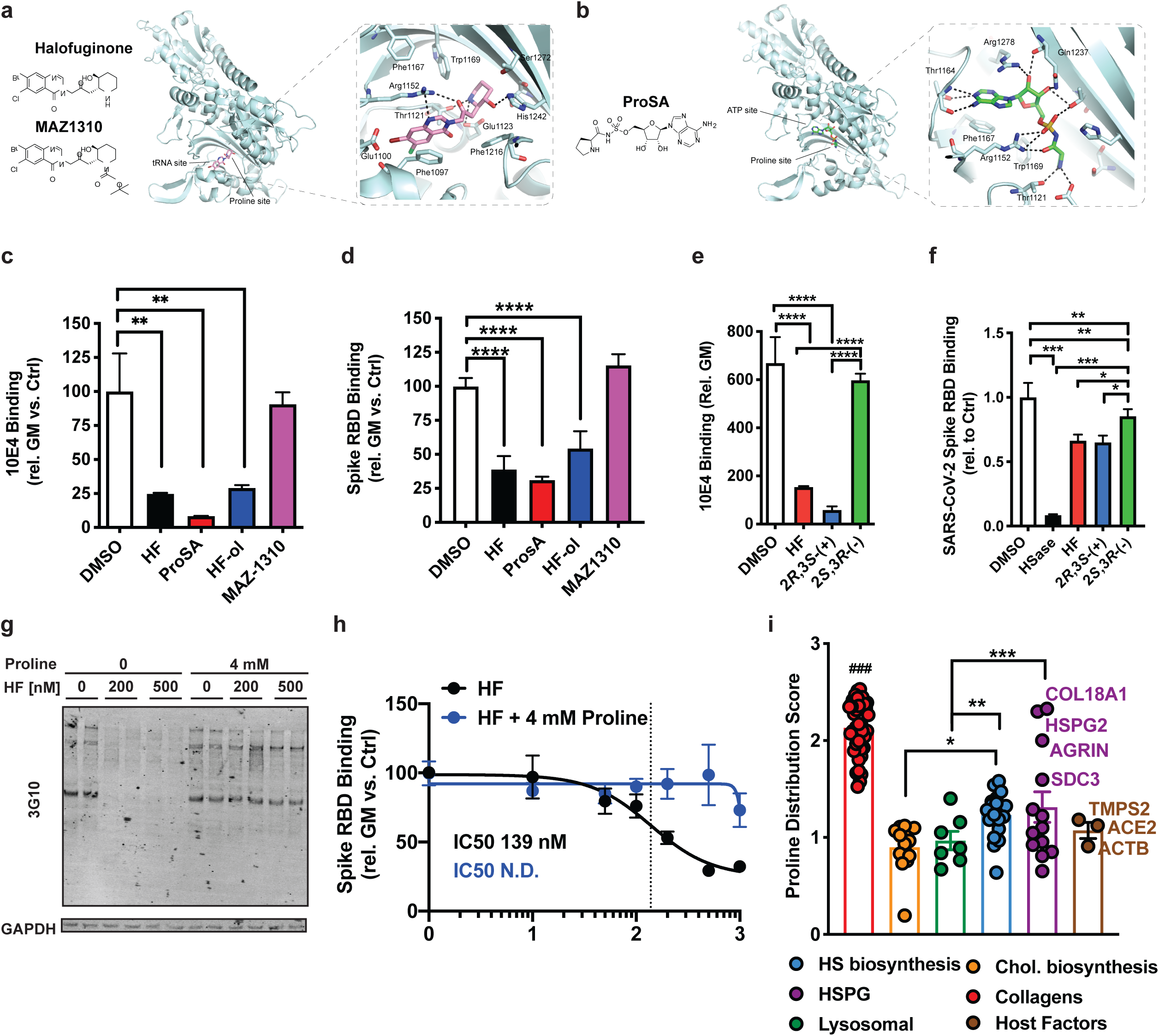

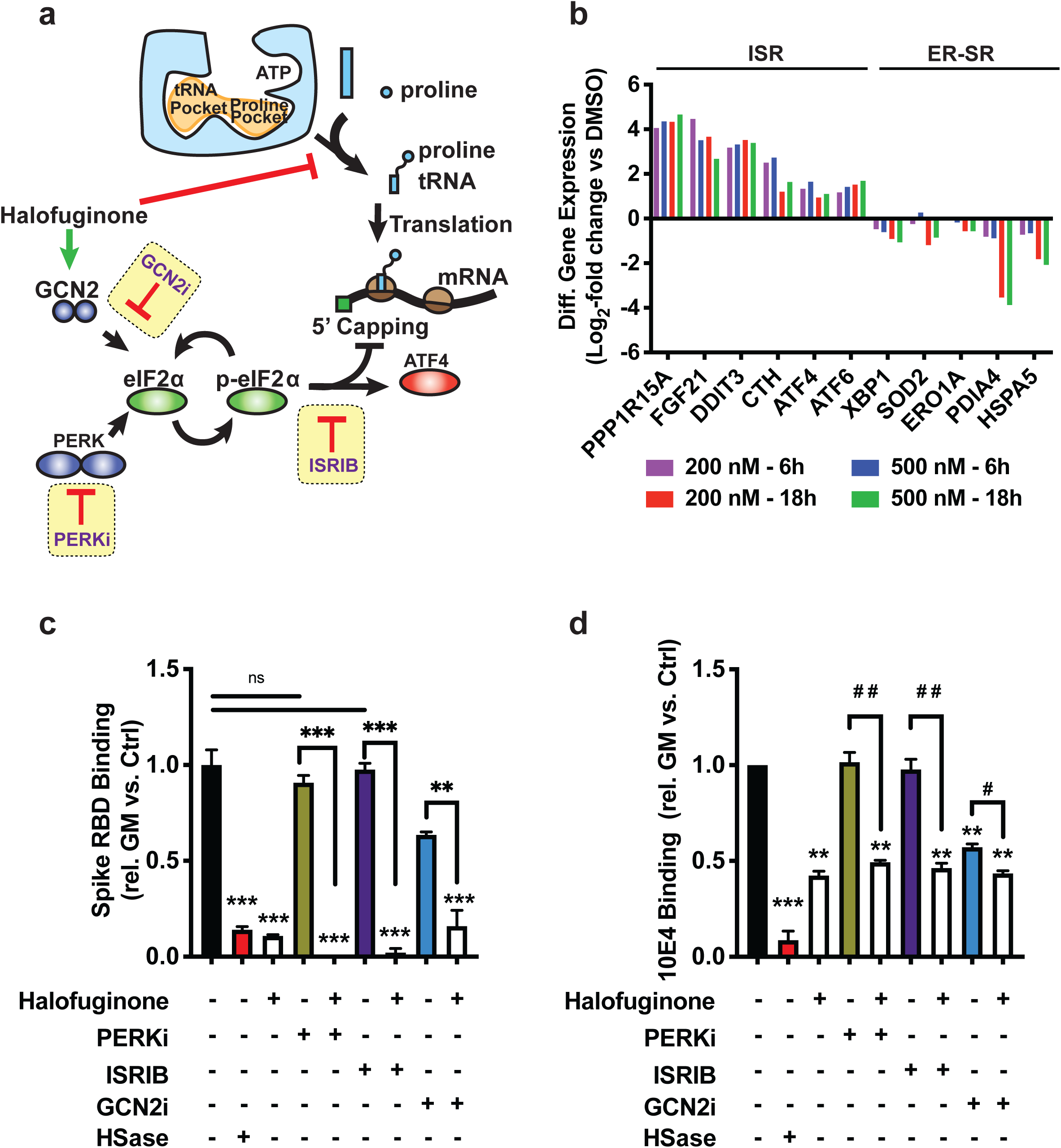

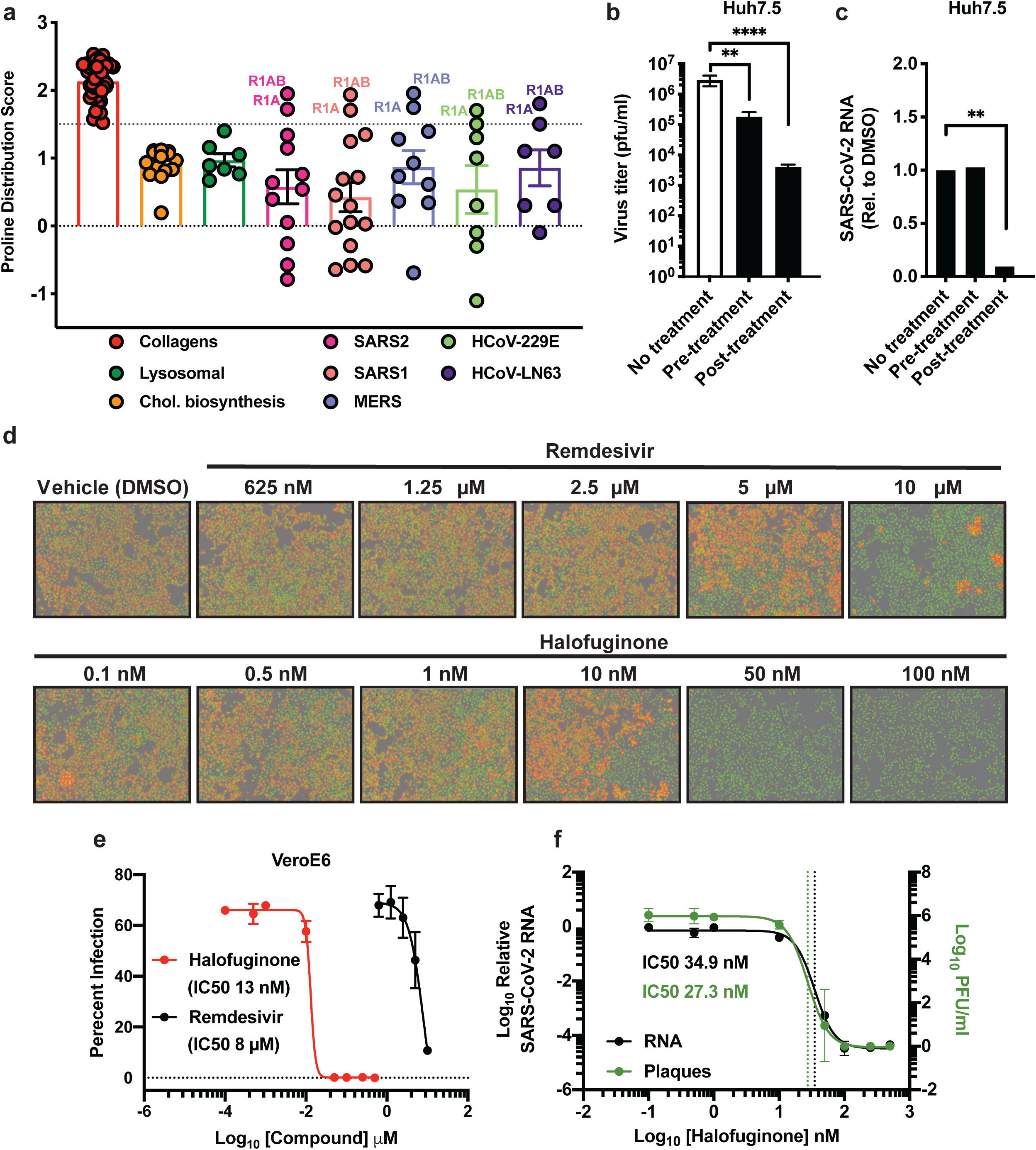

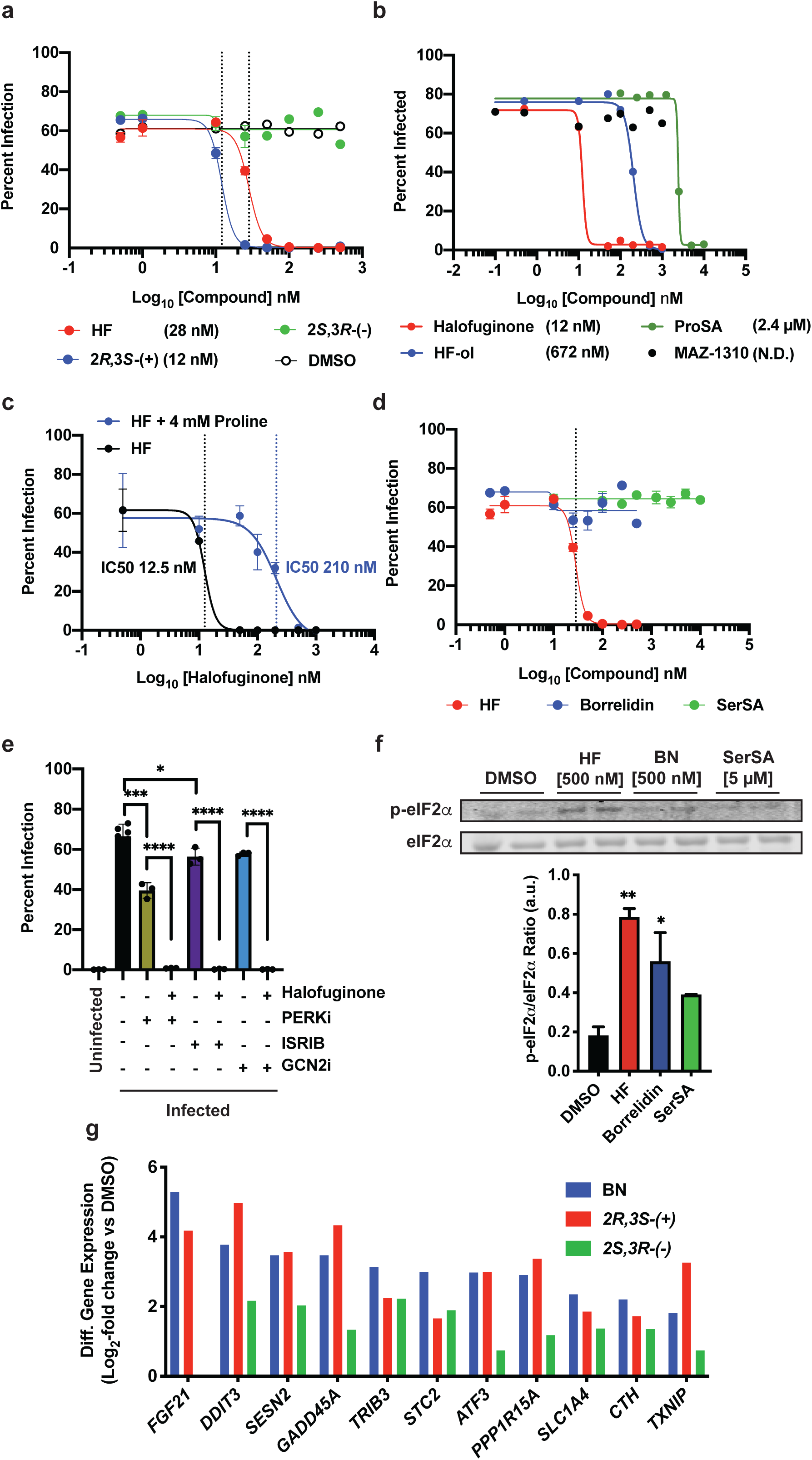

