## Extended Data Figures for "The Prolyl-tRNA Synthetase Inhibitor Halofuginone Inhibits SARS-CoV-2 Infection"

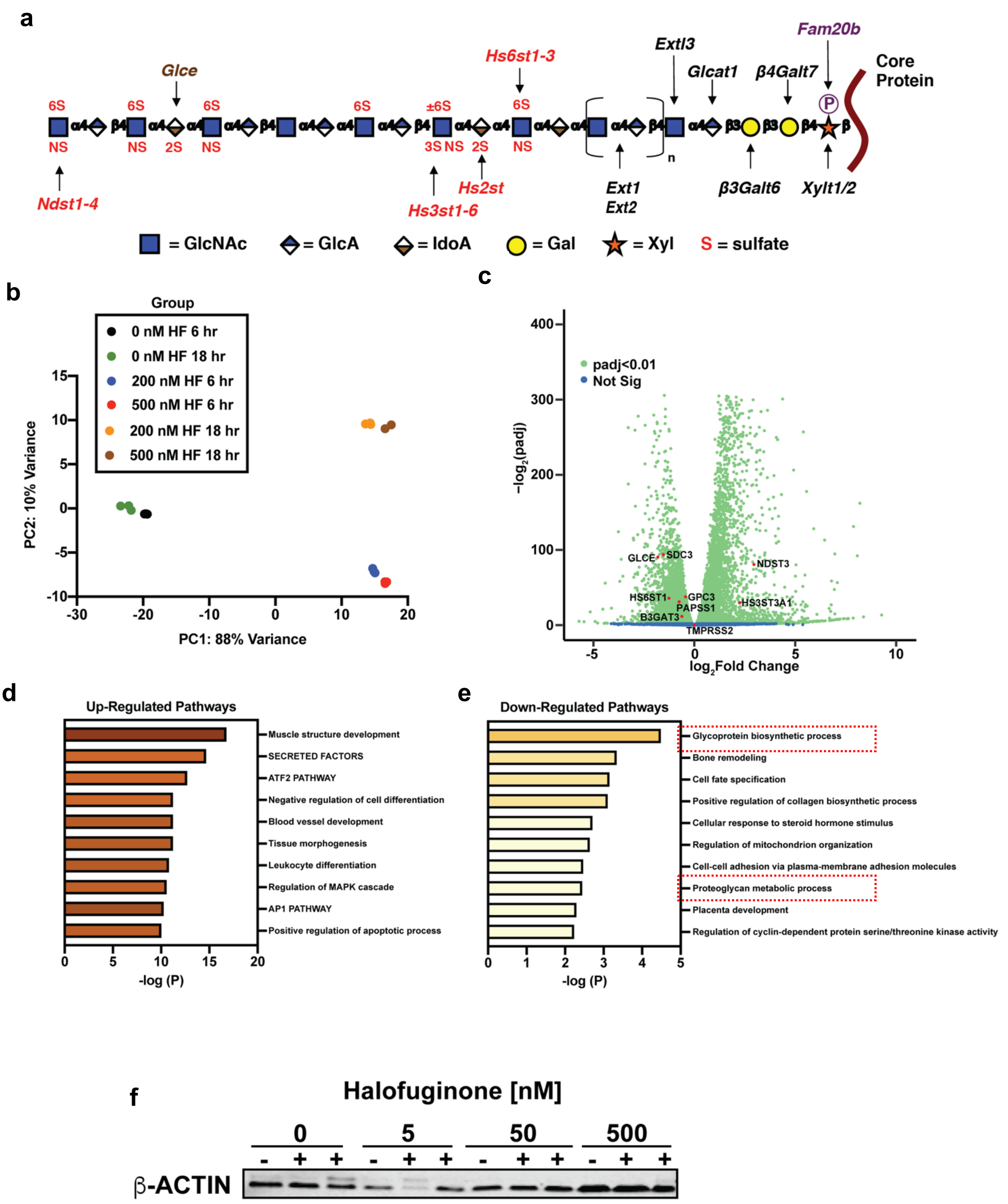


**Supplemental Figure 1 related to Figure 3**

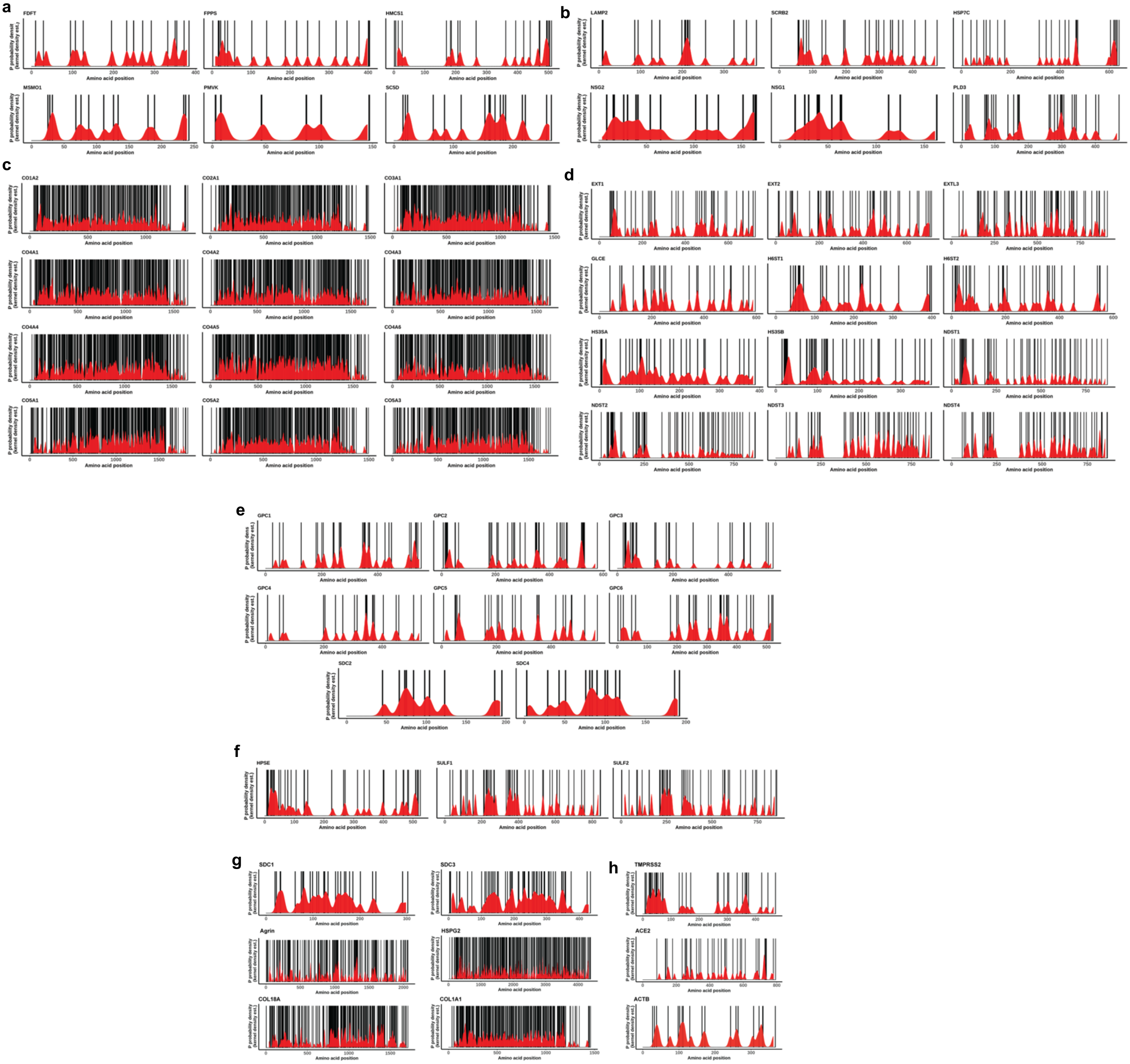


### Supplemental Figure 2 related to Figure 4

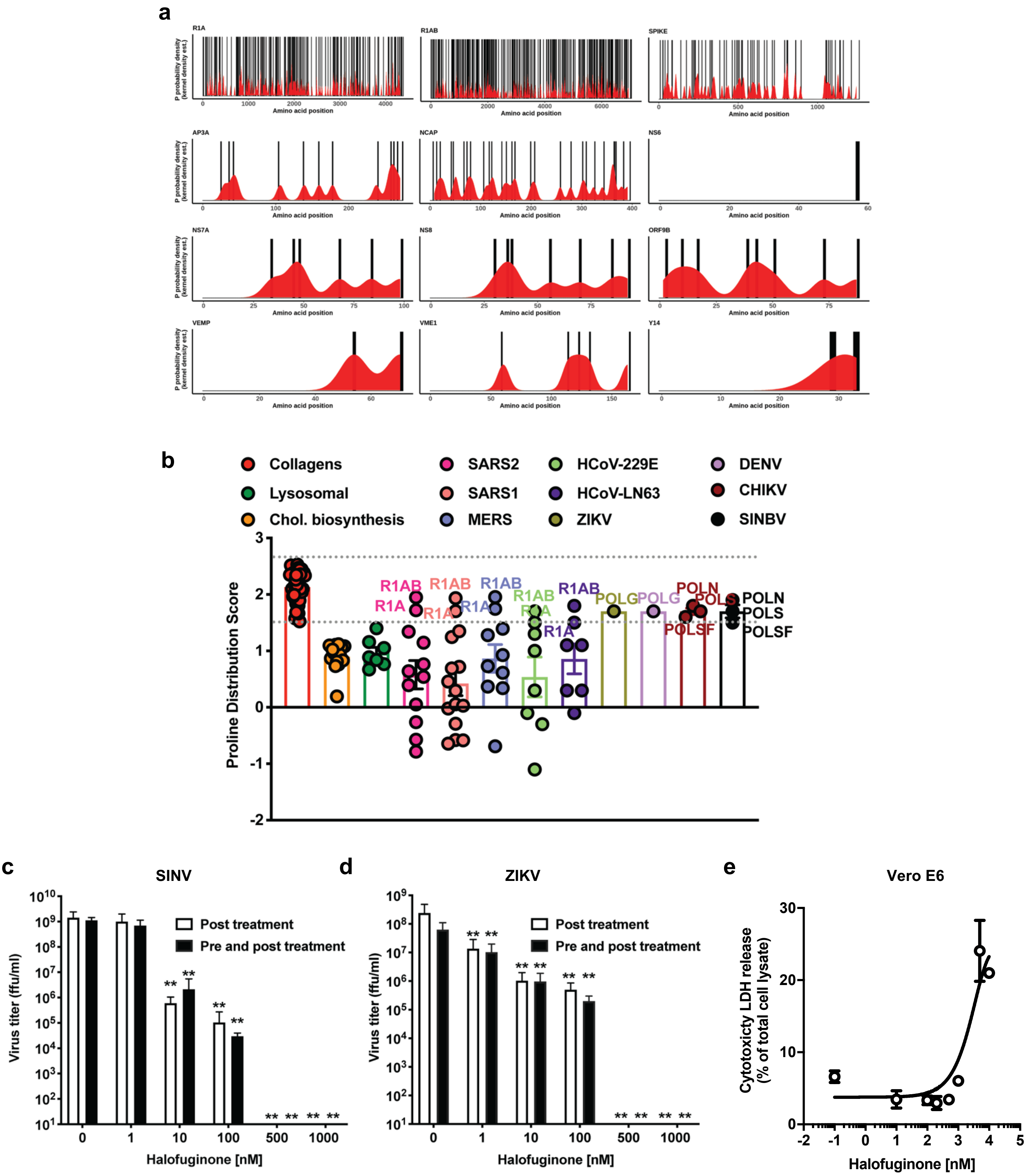


**Supplemental Figure 3 related to Figure 6**

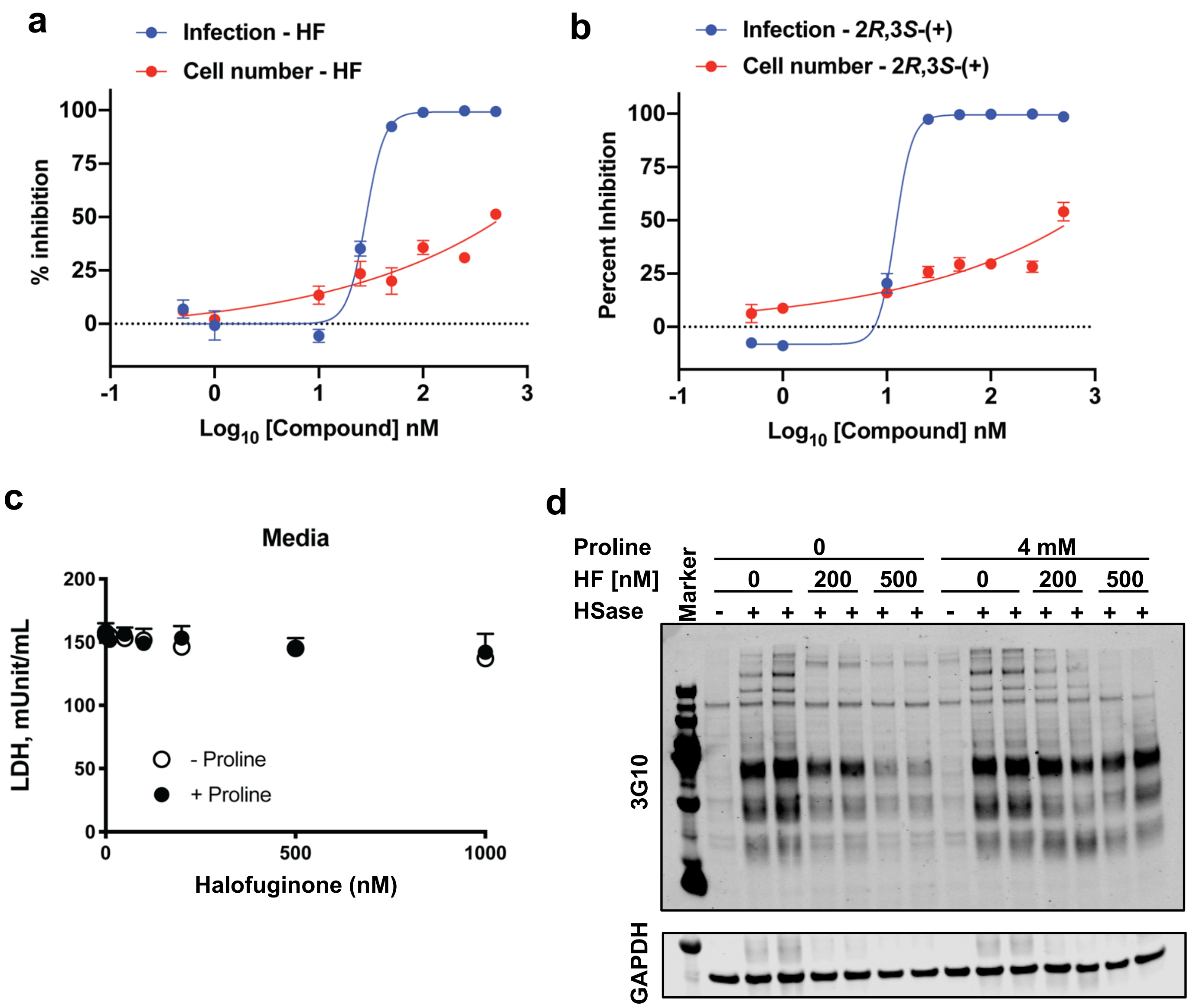


**Supplemental Figure 4 related to Figure 7.**
